# Spatial multi-omics analysis reveals vimentin^high^ macrophages–endothelial cells niche shapes CAFs heterogeneity in colorectal cancer metastasis

**DOI:** 10.64898/2026.08.26.747355

**Authors:** Mingzhou Li, Bingyu Xu, Jieqiong Wu, Zhengyu Zhang, Bingze Chen, Yujie Chen, Danyi Li, Xinrui Tu, Kexin Wang, Zhihan Yang, Yifang Li, Yin Tan, Jinghao Huang, Yunfei Ni, Zilong Chen, Yining Chen, Junfeng Qiu, Sisi Zeng, Li Liang

**Author notes:** **Correspondence: Li Liang,** Department of Pathology, Nanfang Hospital, Southern Medical University, Guangzhou 510515, Guangdong, China. These authors have contributed equally to this work and share first authorship (Mingzhou Li, Bingyu Xu, Jieqiong Wu, Zhengyu Zhang).

## Abstract

The spatial architecture of the tumor microenvironment (TME) is pivotal in the progression of colorectal cancer (CRC) liver metastasis. By applying high-plex spatial multi-omic mapping and neighborhood analysis to a discovery cohort of colorectal cancer primary tumor (PT) and paired liver metastases (LM), we identified a specialized vimentin^high^ macrophages-endothelial cells niche that orchestrates cancer-associated fibroblast (CAF) phenotypes. Mechanistically, in primary tumors, vimentin^high^ macrophages secrete INHBA to activate the ACVR2/TGF-β axis in endothelial cells, driving CAFs toward a myCAF phenotype. Conversely, in liver metastases, these macrophages secrete CXCL9 to trigger the PI3K-Akt/NF-кB/CXCL12 cascade in endothelial cells, directing CAFs toward an iCAF state. Clinically, high niche activity predicts poor survival. Divergent endothelial signaling in primary versus metastatic lesions exposes site-specific stromal vulnerabilities for therapeutic targeting, with architectural features discernible from routine histopathology. These findings reveal a site-specific regulatory mechanism of the macrophage-endothelial niche, offering a novel and clinically significant biomarker for CRC prognosis.

## INTRODUCTION

Colorectal cancer (CRC) ranks as the third most prevalent malignancy worldwide^1^, with approximately 20% of patients presenting with distant metastasis at the time of initial diagnosis^2^. For those with metastatic disease, the 5-year survival rate is dismal, ranging from 12% to 14%^3^. The liver constitutes the predominant site of metastatic spread, affecting roughly 50% of CRC patients over the course of their illness^4,5^. Metastatic progression in CRC is driven not only by tumor cell-intrinsic genetic alterations but also the spatial architecture of the TME^6^. During hepatic dissemination, cellular niches undergo adaptive remodeling, preserving their lineage identity while concurrently acquiring dependencies on the liver microenvironment. Elucidating the molecular determinants of these niche dynamics is therefore essential for understanding metastatic behavior and for improving clinical outcomes in colorectal cancer liver metastasis (CRLM).

Single-cell and spatial transcriptomics have reshaped our understanding of CRC heterogeneity, resolving complex epithelial, immune and stromal ecosystems in both primary tumors and liver metastases^7–9^. Meanwhile, highly multiplexed *in situ* multi-omics platforms, such as multi-omics *in situ* pairwise sequencing (MiP-seq), enable simultaneous detection of DNA, RNA, proteins and other biomolecules at subcellular resolution while preserving native tissue architecture^10,11^. Dissecting the heterogeneity of TME in liver metastases is essential for therapeutic development in CRLM^12^. Across various malignancies, spatial profiling has identified vimentin^high^ macrophages as a distinct immunoregulatory subset driving tumor progression and therapeutic resistance^13^. Tumor-associated endothelial cells (ECs) similarly act as active regulators of CRC progression, governing angiogenesis, immune trafficking and stromal organization^14–17^. Macrophage-centered niches function as integrated spatial units that coordinate matrix remodeling and metastatic niche formation. Nevertheless, the role of vimentin^high^ macrophages in CRC evolution across metastatic sites remains poorly understood, and in particular, whether they collaborate with ECs to establish spatial organized niches that drive stromal adaptation during hepatic colonization is yet to be determined.

Cancer-associated fibroblasts (CAFs) are critical drivers of stromal reorganization during metastatic progression^18^. Single-cell transcriptomic profiling across various solid tumors has resolved functionally distinct CAF states, including myofibroblastic CAFs (myCAFs) that deposit extracellular matrix, inflammatory CAFs (iCAFs) that secrete immunomodulatory cytokines, and antigen-presenting CAF-like states (apCAFs) that regulate T-cell responses^19–21^. In CRC, specific CAF subsets promote desmoplasia, immune exclusion and poor clinical outcomes, with stromal remodeling representing a defining feature of liver metastatic lesions^18,22–26^. Despite this established heterogeneity, the spatial architecture of CAF states across primary and metastatic sites remains unclear. In particular, the local cues governing site-specific activation of myCAF and iCAF program during hepatic colonization have yet to be defined.

In this study, we integrated high-plex spatial multi-omic mapping, spatial transcriptomics, multiplex immunofluorescence and single-cell RNA sequencing on matched primary CRC and their corresponding liver metastases to define the spatial evolution of the TME during hepatic colonization. We uncovered a recurrent vimentin^high^ macrophage–endothelial niche that is conserved across both primary CRC and liver metastases yet displays distinct patterns of CAF organization depending on the tissue location. Spatial neighborhood, cellular niche and topology analyses further revealed that this niche is linked to myCAFs organization in primary tumors, whereas in metastatic lesions it correlates with iCAFs. Integrative transcriptomic and computational analyses support a hierarchical model wherein vimentin^high^ macrophages and endothelial cells cooperate to orchestrate site-specific stromal programs during metastatic adaptation. Finally, by combining spatial validation with deep-learning-based histopathological modeling, we show that the architectural features of this niche are predictive of patient outcome. Taken together, our findings offer a conceptual framework for understanding stromal remodeling in CRLM and identify a candidate microenvironmental vulnerability for precision oncology.

## MATERIALS AND METHODS

### Patient cohorts

This study was approved by the Ethics Committee of Nanfang Hospital, Southern Medical University (Guangzhou, China), in accordance with the Declaration of Helsinki (Approval No. NFEC-2025-481). We analyzed three CRC cohorts. Sex and gender were not considered in the study design, as the research focused on molecular and cellular mechanisms unrelated to sex-based differences. Sex was self-reported for all collected samples.

Cohort 1. CRC liver metastasis was enrolled in the MiP-seq experiments following written informed consent. Paired primary tumor and liver metastasis (LM) samples were collected from each patient.

Cohort 2. To validate the cell cluster distribution patterns identified by MiP-seq, we performed mIHC on paired formalin-fixed paraffin-embedded (FFPE) sections from primary tumors and liver metastases of 20 patients.

Cohort 3. FFPE tissue sections from 560 patients with CRC were stained with H&E, and whole-slide images were acquired for HE2Surv model development.

Cohort 4. FFPE tissue sections from 56 patients with CRC form Zhujiang Hospital, Southern Medical University (Guangzhou, China) were stained with H&E, whole-slide images and prognostic information were acquired for HE2Surv model validation.

Cohort 5. FFPE tissue sections from 122 patients with CRC form the Fourth Affiliated Hospital of Hebei Medical University were stained with H&E, whole-slide images and prognostic information were acquired for HE2Surv model validation.

Cohort 6. FFPE tissue sections from 26 patients with CRC form the Guangdong Provincial People’s Hospital were stained with H&E, whole-slide images and prognostic information were acquired for HE2Surv model validation.

### Mouse models and animal studies

C57BL/6 mice were obtained from the Guangdong Medical Laboratory Animal Center (Guangzhou, China). Animals aged 4–7 weeks were housed in specific pathogen-free (SPF) facilities at Southern Medical University under controlled temperature (22 ± 2°C) and humidity (50–60%) with a 12-hour light/dark cycle. All animal experiments were performed in accordance with the ARRIVE guidelines and approved by the Institutional Animal Care and Use Committee (IACUC) of Southern Medical University.

To enrich for colorectal cancer cells with enhanced metastatic capacity, 2 × 10⁵ MC38 cells were intrasplenically injected into C57BL/6 mice to establish experimental liver metastases. Following tumor development, hepatic metastatic foci were excised, dissociated into single-cell suspensions, and expanded in vitro. These cells were subsequently re-injected into the spleens of recipient mice. This in vivo selection cycle was repeated for six sequential passages to generate the high-metastatic MC38 (MC38-HM) subline. For the orthotopic model, mice underwent midline laparotomy, and the cecum was gently exteriorized onto sterile saline-moistened gauze. 1×10⁶ MC38 or MC38-HM cells in 25 μL of 100% high concentration Matrigel (Corning) were implanted into the cecum using a 30-gauge needle (Braun, Melsungen, Germany). The cecum was returned to the peritoneal cavity, and the abdominal wall was closed in layers and the incision was closed.

### Cell culture

MC38 (mouse colon adenocarcinoma), HUVEC (human umbilical vein endothelial cells), and THP-1 (human monocytic leukemia) were obtained from the American Type Culture Collection (ATCC). THP-1 cells were maintained in RPMI 1640 medium (Gibco), whereas MC38, MC38-HM, and HUVEC cells were cultured in DMEM (Gibco). All media were supplemented with 10% fetal bovine serum (FBS) and 1% penicillin-streptomycin. Cells were incubated at 37°C in a humidified atmosphere containing 5% CO₂. All cell lines were routinely tested for mycoplasma contamination and confirmed to be negative.

### Multiplex immunohistochemistry

Four-color multiplex immunohistochemistry (mIHC) was performed as previously described^27^. Primary antibodies against CD31 (CST, 77699), CD68 (Proteintech, 66231-2-Ig), and Vimentin (Abcam, ab92547) were sequentially applied. Image acquisition, spectral unmixing, and signal quantification were performed using an LSM 880 confocal microscope (Zeiss) or the Vectra Polaris imaging system (Akoya Biosciences).

The Seven-color mIHC staining was performed by Guangzhou RuiGene Biotechnology Co., Ltd. mIHC staining was obtained using PANO 7-plex IHC kit (Panovue, Beijing, China,cat no.0004100100). Tissue sections were sequentially incubated with primary antibodies against α-SMA (Abcam, ab5694), FAP (Abcam, ab314456), IL-6 (Abcam, ab9324), CD31 (CST, 3528), CD68 (CST, 76437), and Vimentin (CST, 5741), followed by horseradish peroxidase-conjugated secondary antibody and tyramide signal amplification (TSA). Antigen retrieval was performed by microwave heating after each TSA cycle. Upon completion of all antigen labeling, nuclei were counterstained with 4′,6-diamidino-2-phenylindole (DAPI; Sigma-Aldrich).

### Hematoxylin and eosin staining

Formalin-fixed, paraffin-embedded tissue sections were baked at 65°C for 1 hour, deparaffinized in xylene, and rehydrated through a graded ethanol series. Sections were stained with hematoxylin for 5 min, differentiated in 1% acid alcohol, and rinsed in running tap water until optimal blueing. Following eosin counterstaining, sections were dehydrated through graded ethanols, cleared in xylene, and mounted with neutral mounting medium. Whole-slide images (WSIs) were acquired using a Aperio AT2 at 20×.

### qRT-PCR and RNA sequencing

Total RNA was extracted using TRNzol Universal Reagent (AG21102) and reverse-transcribed using the PrimeScript RT Reagent Kit (Takara, RR037Q). Quantitative real-time PCR was performed with SYBR Green Master Mix (Vazyme, Q711-02) on an ABI Prism 7500 Sequence Detection System (Applied Biosystems). Relative gene expression was normalized to β-actin and quantified using the 2^(−ΔΔCt) method. Primers were designed using Primer Express software (Applied Biosystems), and sequences were listed in Supplementary Table 1. All experiments were performed in triplicate with three biological replicates.

Total RNA was extracted from primary MC38 tumors and paired primary and liver metastatic MC38-HM lesions using TRIzol reagent (Invitrogen). RNA sequencing was performed as previously described ^28^.

### Enzyme-linked immunosorbent assay (ELISA)

TGF-β1 and CXCL12 levels in culture supernatants of HUVEC cells were measured using human TGF-β1-specific ELISA kits (Elabscience, E-EL-0162) and human CXCL12-specific ELISA kits (Elabscience, E-EL-H0052) according to the manufacturer’s instructions. All experiments were performed in triplicate with three biological replicates.

### MiP-seq probe design

The MiP-seq probe set comprised four components: (1) padlock probe pairs for target recognition; (2) an RCA initiator primer for target-dependent rolling-circle amplification; (3) fluorophore-conjugated detection probes for amplified product visualization; and (4) anchor and query probes for *in situ* sequencing. Design criteria were as follows: target sequences were computationally screened for specificity using an in-house algorithm; exonic regions proximal to the 5′ terminus were prioritized with two independent sites selected per gene; padlock probes contained 12–16 bp target-complementary arms at both termini flanking a central backbone with anchor primer-binding and dual-barcode sequences; the RCA initiator primer contained a 5′ target-complementary arm (12–16 bp) and a 3′ sequence annealing to the padlock backbone; detection probes (CY3-P4, 488-1, CY3-1, CY5-1, and CY7-1) enabled RCA efficiency assessment and multiplex expression profiling; *in situ* sequencing probes included three anchor primers (anchor-1 through anchor-3) for sequential hybridization and four query probes (seq-1–seq-4) conjugated to Alexa Fluor 488, 546, 594, and 647, respectively. For paired-end sequencing, three anchor primers were employed; for pairwise sequencing, three probe pairs were used. All oligonucleotides were synthesized by Sangon Biotech (Shanghai, China); sequences were listed in Supplementary Table 2.

### Co-detection of protein and mRNA by MiP-seq

Protein targets were detected using antibody–oligonucleotide conjugates (5 ng µl⁻¹ per antibody). For simultaneous detection of 30 immune-related proteins, listed in Supplementary Table 3, and 18 marker gene transcripts in human CRC, 3 µm sections were cut from FFPE tissue blocks.

FFPE section preparation. Sections were baked at 60 °C for 1 h, dewaxed twice in Histo-Clear II (National Diagnostics, HS-202) for 5 min each, then in 10% ethanol/90% Histo-Clear II for 5 min. Rehydration was performed through a graded ethanol series (100%, 95%, 85%, 70%, 50%, 30%; 5 min each), followed by washes in DEPC-treated water and PBS (5 min each). Antigen retrieval was performed with proteinase K (10 µg ml⁻¹ in PBS) at 37 °C for 10 min, followed by fixation with 4% paraformaldehyde for 10 min. Sections were dehydrated through a reverse ethanol series (30–100%, 1 min each), washed twice in PBS (5 min each), and blocked with 2% BSA and 5% goat serum in PBS for 1 h.

Antibody staining and MiP-seq. The BD AbSeq Immune Discovery Panel (30 antibodies, BD Biosciences, 625970) was applied at 1:5,000 dilution for 1 h at room temperature. Sections were then processed through the standard MiP-seq workflow: probe hybridization, ligation, rolling-circle amplification, and signal decoding as previously described^10^. For protein–RNA co-detection, an additional T4 DNA ligase step was inserted following the SplintR ligation to ensure efficient circularization of padlock probes bound to antibody-conjugated oligonucleotides. The MiP-seq was performed using commercial kits (Kunyu Biotech, Wuhan, China) according to the manufacturer’s protocol.

### Raw image processing, cell segmentation and quality control of MiP-seq

After obtaining the raw images, we performed image registration using BigWarp in ImageJ to eliminate image rotation errors caused by extraneous factors, thereby enhancing image reliability. Bigwarp is a deformable image alignment tool based on a Java implementation of the Thin Plate Spline (TPS) model. Following image registration, we utilized the RS-FISH plugin in ImageJ for fluorescent signal quantification and localization. RS-FISH maintains high detection accuracy and low localization error across a broad range of signal-to-noise ratios. Subsequently, we employed Cellpose for cell segmentation to delineate the boundaries of individual cells. After excluding low-quality cells and background noise, a total of 587,513 high-quality cells were segmented. The spatial coordinates of each cell, along with the measured quantities of various proteins and genes, were compiled into a table for downstream analysis.

### Cell type definition

We utilized 15 commonly used markers to annotate all cells. The entire cell population was classified into 13 distinct cell types, encompassing immune cells, epithelial cells, endothelial cells, and stromal cells. We employed heatmaps to visualize the annotation results for each cell, demonstrating the accuracy and robustness of our cell annotation.

### Quantification of pairwise co-occurrence between cellular neighborhoods

We defined the cellular neighborhood of each cell as the 8 nearest neighboring cells. We constructed a spatial connectivity graph of cell types using the k-nearest neighbors (kNN) algorithm and calculated z-scores using a random permutation algorithm. The z-scores were employed for standardization to eliminate potential inter-sample variations, quantitatively representing the level of attraction or repulsion between cells. The co-occurrence coefficients of identical cell pairs from colorectal cancer PT and liver metastasis LM groups were then linearly summed and averaged, respectively, to represent the degree of co-occurrence between different cell types in both groups. We compared identical cell pairs across different samples between the two groups, visualized the co-occurrence degrees of identical cell pairs using boxplots, and performed significance analysis using paired Wilcoxon test and Wilcoxon rank-sum test.

### Definition of cellular niches

We again defined the neighborhood of each cell as the 8 nearest neighboring cells. We first performed cell type alignment across all samples. Subsequently, we calculated the abundance of different cell types surrounding each cell and constructed a cellular neighborhood matrix. We then applied K-means clustering to the cellular neighborhood matrix, partitioning the neighborhoods into 10 distinct niches. Finally, we visualized the cellular composition of each niche using stacked bar charts, with particular emphasis on the proportions of iCAFs, myCAFs, VIM^+^ macrophages, and vascular endothelial cells.

### Construction of Voronoi Diagrams and Shared-Edge Analysis

Real cell boundaries are often unsuitable for shared-edge analysis due to the selection of cell segmentation algorithms. Therefore, we employed Voronoi tessellation to calculate shared-edge relationships between cell distributions. We utilized the “voronoi_finite_polygons_2d” function to address the distortion of peripheral points inherent in conventional Voronoi tessellation by generating virtual vertices and sorting polygon vertices in counterclockwise order. Subsequently, we quantified the number and proportion of shared edges between different cell types within the Voronoi diagram and visualized these results using heatmaps.

### Processing of Visium Spatial Transcriptomics Data and colocalization analysis

We downloaded Visium spatial transcriptomics data of colorectal cancer primary tumors and liver metastases from the GEO database. We employed the Robust Cell Type Decomposition (RCTD) algorithm to perform deconvolution and cell annotation of the spatial transcriptomics data, utilizing the quality-controlled and manually annotated single-cell RNA-seq data from GSE178318, comprising colorectal cancer primary tumors and matched liver metastases, as the reference single-cell dataset. We assumed that each spot contained 15 cells. Subsequently, we visualized the spatial distribution of myCAFs, iCAFs, VIM^+^ macrophages, and vascular endothelial cells, and selected specific samples for colocalization analysis. We plotted the proportions of two target cell types in each Visium spot on the x- and y-axes, fitted trend lines, and conducted significance analysis.

### Cellular Neighborhood Analysis of Visium Spatial Transcriptomics Data

Following RCTD deconvolution, we defined the 6 surrounding spots of each spot as its cellular neighborhood and constructed a kNN-based cell proximity matrix. Subsequently, we calculated z-score indices between different cell pairs through random permutation testing to quantify the degree of cellular co-occurrence. We visualized these results using boxplots and assessed statistical significance using paired Wilcoxon tests.

### Processing of scRNA-seq Data

We obtained and integrated single-cell sequencing datasets from the GEO database, including normal colorectal mucosa, invasive margin of colorectal cancer, non-metastatic colorectal cancer tissues, primary tumor tissues of metastatic colorectal cancer, liver metastatic tissues of metastatic colorectal cancer, and normal liver tissues (GSE132465, GSE144735, GSE298084, GSE245552, GSE221575, GSE302903), totaling 113 samples; as well as anti-PD-1 ICB therapy-resistant colorectal cancer tissues (GSE236581) from 22 patients; and additionally, paired patient-derived normal colorectal tissues, primary metastatic lesions, liver metastases, and normal liver tissues (GSE178318, GSE225857) from 13 patients. Cell clustering and manual cell annotation were performed separately for each dataset, and the proportions of cell types were visualized using stacked bar charts.

We analyzed the above integrated single-cell datasets separately using the Seurat package (version 5.5.1). For quality control, cells with gene counts less than 300 or greater than 10,000, mitochondrial percentage greater than 20%, or total RNA counts less than 500 were identified as contaminated and excluded. For doublet removal, cells that expressed two or more mutually exclusive markers after clustering were identified as doublets and excluded. For batch effects, we utilized the Harmony (version 2.0.5) package in R to remove batch effects. We subsequently annotated cells into 8 distinct cell types based on commonly used marker genes, including lymphocytes, endothelial cells, epithelial cells, and myeloid cells. We then performed further clustering on myeloid cells, yielding 7 distinct subtypes. We further clustered fibroblasts into 3 distinct subtypes according to established conventions. Subsequently, we employed the CellChat (version 2.2.0) package to analyze intercellular communication intensity. We utilized the Monocle3 (version 1.4.25) package to infer pseudotemporal relationships in colorectal cancer PT and LM groups.

### Prognostic Analysis of Cellular Niches

We first downloaded bulk RNA-seq data and paired survival prognostic information from 339 patients in the TCGA database. We subsequently calculated characteristic genes for VIM⁺ macrophages, iCAFs and myCAFs, selecting the top 30 genes based on log fold change (log FC) plus 2 characteristic marker genes as the signature gene set for each cell type. The signature gene set were listed in Supplementary Table 4. We then computed ssGSEA scores for each patient, stratified patients into high- and low-score groups, calculated hazard ratios (HR), and generated Kaplan-Meier (KM) survival curves to quantify the impact of niche composition on patient prognosis.

### Analysis of Multiplex Immunofluorescence Data

QuPath software was utilized for the analysis of mIHC images. Following import of all raw mIHC images into QuPath, we performed cell segmentation within regions of interest (ROIs) and exported the coordinates of each cell along with fluorescence intensities across all channels. After quality control exclusion of low-quality cells, we annotated cells into 5 distinct types based on these data: iCAFs, myCAFs, vascular endothelial cells, VIM^+^ macrophages, and VIM^−^ macrophages. We subsequently defined a 40-μm radius surrounding each vascular endothelial cell as its cellular neighborhood, calculated the distances from each vascular endothelial cell to its nearest cells of various types, and computed the mean values. We then compared the average distances between vascular endothelial cells and myCAFs or iCAFs across different groups, and validated statistical significance using paired Wilcoxon tests.

### Online database analysis

Bioinformatic analyses, including immune infiltration analysis and survival analysis, were conducted using the BEST (Biomarker Exploration for Solid Tumors) online platform (https://rookieutopia.com/)^29^. BEST is a publicly accessible web server that integrates multi-omics datasets from over 10,000 solid tumor samples across 27 cancer types, with transcriptome data uniformly re-annotated based on the GRCh38 patch 13 reference genome. For immune infiltration analysis, the abundance of infiltrating immune cells was estimated using ssGSEA. For survival analysis, Kaplan–Meier curves were generated, and the log-rank test was used to compare survival differences between high- and low-expression groups.

### Construction of the HE2Surv Model

We constructed the HE2Surv model using whole slide images (WSIs) of HE-stained sections from colorectal cancer patients at our center. After excluding deaths from non - cancer causes, we obtained 560 WSIs with accompanying prognostic information for HE2Surv model construction, and additionally included 56, 122, and 26 WSIs with prognostic information from the Zhujiang, HeBei, and GDPH cohorts, respectively, for external validation of the model. The HE2Surv model comprises two components: Universal Non - locked Image encoder version 2 (UNI v2) and Clustering - constrained Attention Multiple instance learning (CLAM). Each WSI was partitioned into 36 tiles. Following exclusion of blank background regions, the tiles were processed by pre - trained UNI v2 for unbiased extraction of histopathological features. UNI v2 extracted 1,536 features from each tile. These features were subsequently employed to train the CLAM model. To enhance training stability, we adopted a five - fold cross - validation approach for prognostic model development. We then designated 5 - year survival as the critical clinical endpoint, generated receiver operating characteristic (ROC) curves, and calculated the area under the curve (AUC).

### Visualization of HE2Surv Attention Maps

CLAM is an attention-based deep learning model. To elucidate which histopathological features HE2Surv relies upon for accurate prognostic prediction, we transformed its attention weights into heatmaps following z-score standardization, with red indicating high attention and blue indicating low attention. We adjusted the heatmap transparency to 0.6 and overlaid it onto the original H&E images to intuitively display regions of high model attention.

### Quantification and statistical analysis

Data were representative of three or more independent experiments. Statistical analyses were performed using GraphPad Prism 9 and SPSS 20. Differences between two groups were assessed using two-tailed unpaired t tests. For multiple comparisons, one-way or two-way analysis of variance (ANOVA) was applied with Tukey correction for post hoc adjustment. Tumor growth curves was analyzed using a mixed-effects model with two-way ANOVA followed by Tukey’s post hoc test. *p < 0.05, **p < 0.01, ***p < 0.001, ns = not significant.

## RESULTS

### Integrative spatial and transcriptomic profiling identifies vimentin^high^ macrophages as a critical driver of CRLM and immunotherapy resistance

To characterize the spatial heterogeneity and evolutionary dynamics of CRC during hepatic colonization, we applied high-plex, high-resolution spatial multi-omic mapping using MiP-seq to a cohort of 20 paired primary CRC and liver metastasis samples derived from 5 patients. To validate the robustness of our spatial mapping, we additionally analyzed an independent validation cohort comprising 28 paired primary and metastastic samples from 15 patients using mIHC coupled with intercellular distance quantification (Fig. 1a). Our tailored panel, which included 30 proteins and 18 genes, enabled simultaneous visualization of major cellular lineages, including intestinal epithelial cells, immune cells, fibroblasts, vascular and lymphatic endothelial cells (Fig. 1b,c). Following stringent cell segmentation and quality control, we assembled a comprehensive spatial atlas encompassing 587,513 cells, which were subsequently subjected to cellular neighborhood and niche analyses. *In situ* imaging of the 20 paired samples from the 5 patients is presented in Fig. 1d.

**Figure 1.**
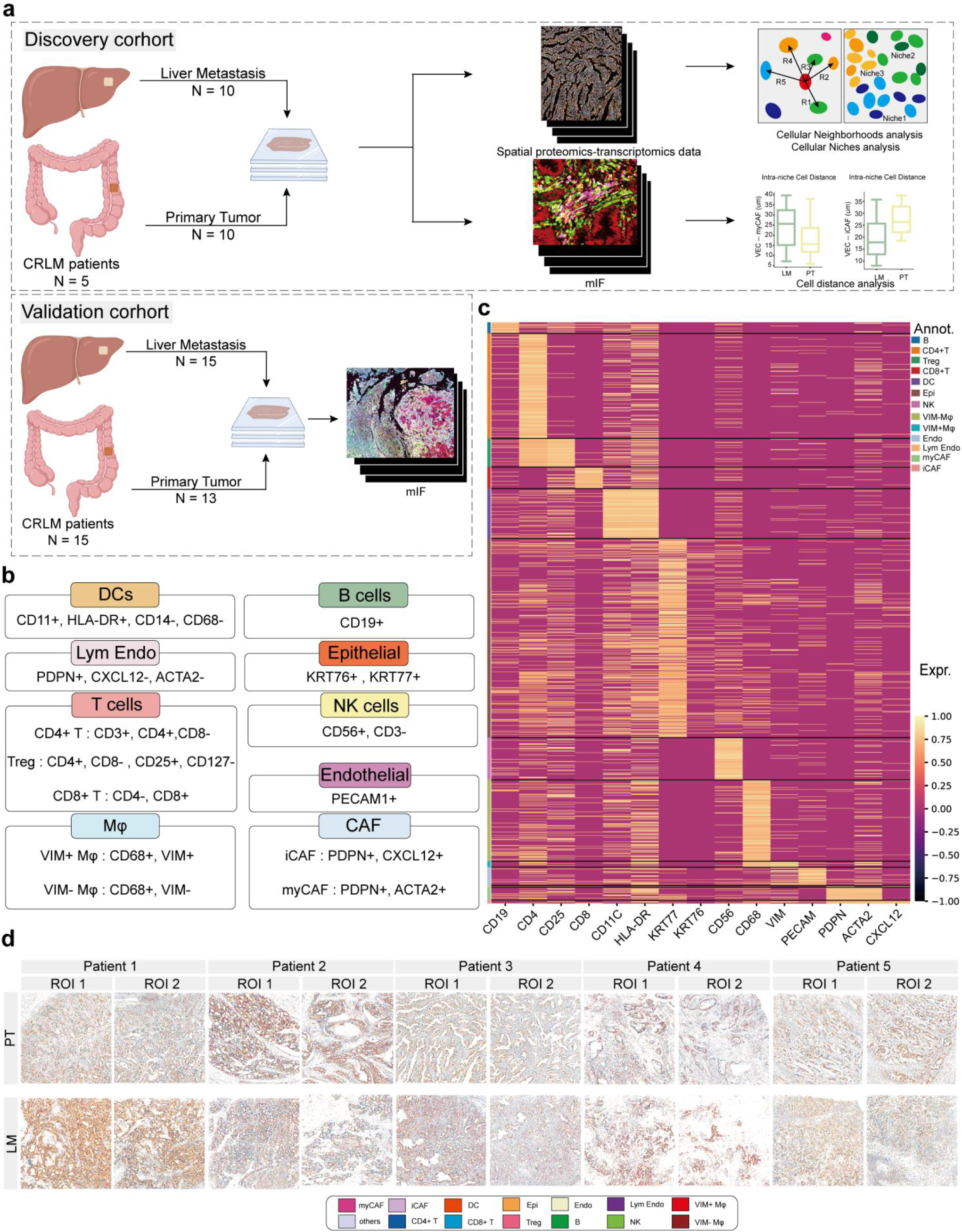
High-plex spatial multi-omic mapping delineates the cellular architecture of primary colorectal cancer and paired liver metastases. (a) Schematic of the study design. Overview of the experimental workflow, including spatial multi-omics (MiP-seq, discovery cohort, n=5 patients, 20 paired PT and LM samples) and multiplex immunofluorescence (mIHC, validation cohort, n=15 patients, 28 samples) integrated with downstream cellular neighborhood, cellular niche, and inter-cellular distance analyses. (b) Panel design and marker signatures. Selection of 30 protein and 18 transcript markers used for cell-type definition, distinguishing immune cells, epithelial cells, endothelial cells, and distinct macrophage and CAF subpopulations. (c) Validation of cellular annotation. Heatmap exhibiting the expression profiles of key marker proteins and genes across 13 annotated major cell lineages among 587,513 segmented single cells. (d) Representative spatial cell-type maps. Spatial distribution maps of annotated cellular populations across 20 ROI pairs from 5 matched CRC patients (PT, top; LM, bottom).

To delineate the evolutionary trajectory of CRC from initiation through progression to liver metastasis, we constructed a single-cell sequencing dataset by integrating publicly available scRNA-seq datasets encompassing normal colorectal mucosa, normal liver tissue, colorectal cancer tumor margins, non-metastatic CRC, primary lesions of metastatic CRC, and matched liver metastases (Supplementary Fig. 1a-c). We found that vimentin expression was independent of other macrophage markers (Supplementary Fig. 1d). Notably, the proportion of vimentin^high^ macrophages exhibited a stepwise increase along the continuum of CRC progression and metastasis (Supplementary Fig. 1e,f). This observation was further corroborated by multiple bulk RNA-seq datasets, which demonstrated that vimentin^high^ macrophages were preferentially enriched in tumor tissues, and that their abundance was tightly correlated with CRC progression and metastatic evolution, significantly correlating with worse patient prognosis (Supplementary Fig. 1g-m).

To explore the therapeutic relevance of this population, we incorporated single-cell RNA sequencing data from CRC patients receiving immunotherapy, stratified by clinical response-including stable disease (SD), partial response (PR), and complete response (CR). We found that the proportion of vimentin^high^ macrophages was highest in the SD group, and declined progressively with improving treatment response. This pattern was further validated in multiple large-scale bulk RNA-seq cohorts of CRC immunotherapy, where the abundance of vimentin^high^ macrophages effectively predicted immunotherapeutic outcomes (Supplementary Fig. 2a-i).

### Vimentin^high^ macrophage–EC niche may direct CAFs subtype specification via spatial proximity in CRLM

To delineate the spatial architecture of TME, we performed spatial neighborhood analysis and utilized heatmaps to visualize cellular proximity, from which potential intercellular interactions were inferred (Fig. 2a). This analysis revealed that vimentin^high^ macrophages consistently interacted with ECs in both PT and matched LM. Notably, ECs displayed site-dependent interaction preferences with distinct CAF subtypes: they were closely associated with myCAFs in PT samples, whereas they preferentially interacted with iCAFs in LM samples (Fig. 2b,c). To further assess the specificity of the vimentin^high^ macrophage-EC niche, we found that vimentin^low^ macrophages did not exhibit significant interactions with ECs (Fig. 2c). These findings were further corroborated by cellular niche analysis (Fig. 2d,e). By clustering the PT and LM landscapes into 10 distinct cell niches (CNs) each, we identified that vimentin^high^ macrophages, ECs, and their corresponding CAF subtypes were coordinately enriched within CN5 and CN6 in PT samples and within CN3 of LM samples, while remaining concurrently depleted in all other niches (Fig. 2f). Collectively, these data suggest that vimentin^high^ macrophages and ECs assemble into a specialized vimentin^high^ macrophage-EC niche, that is closely associated with myCAFs in primary lesions but transitions to iCAFs in metastatic lesions, thereby contributing to the evolutionary dynamics of CRLM microenvironment.

**Figure 2.**
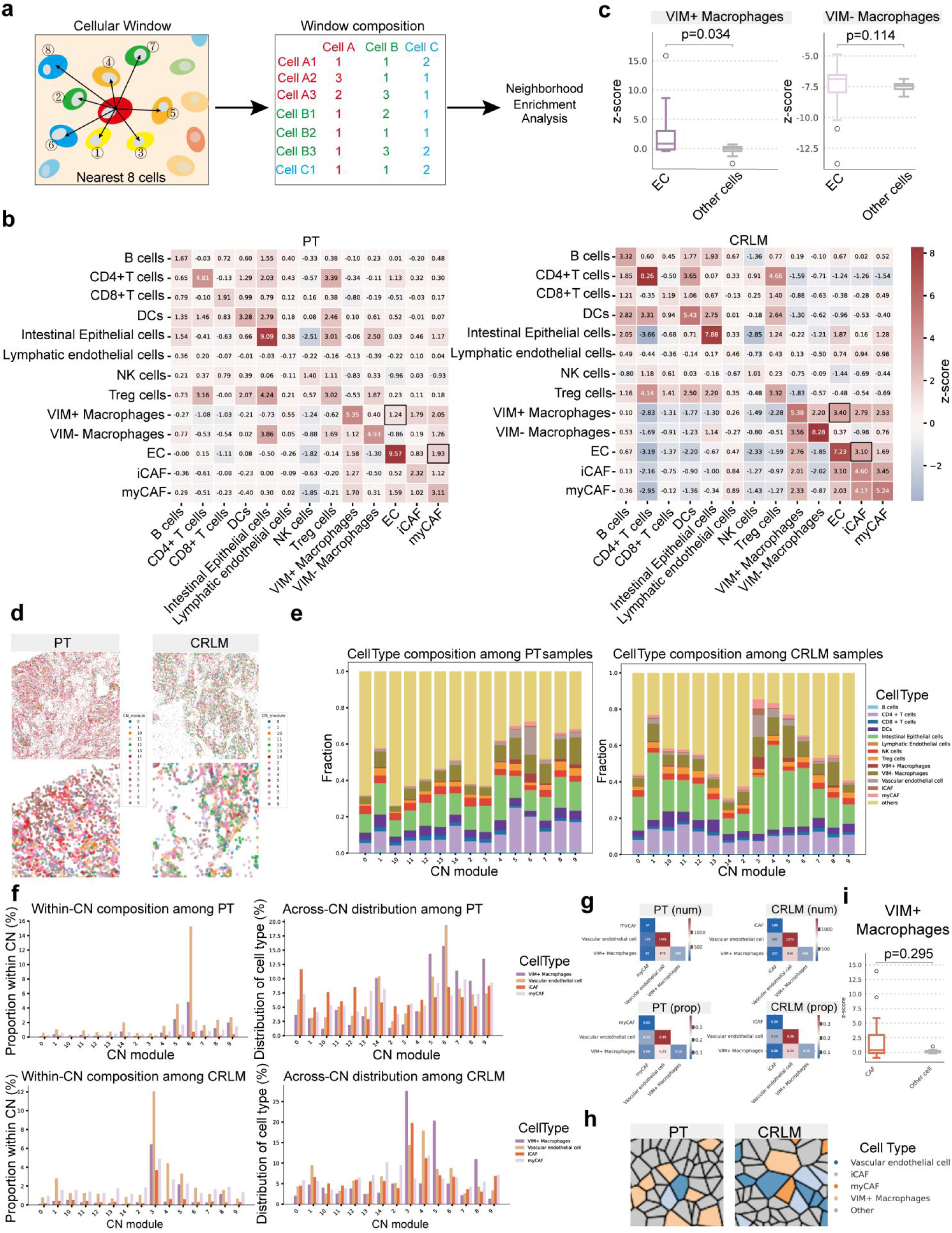
Identification of a specialized vimentin^high^ macrophage–endothelial cell niche associated with site-specific CAF phenotypes. (a) Workflow of spatial neighborhood analysis. Schematic illustrating the 8-nearest-neighbors (kNN) cellular window approach used for spatial proximity quantification. (b) Cellular interaction heatmaps. Z-score enrichment matrices depicting spatial co-occurrence or avoidance among cell types in primary tumors (PT, left) and paired liver metastases (CRLM, right). Black boxes highlight interactions within the macrophage–EC–CAF axis. (c) Specificity of macrophage–EC interactions. Comparison of spatial proximity (z-score) between ECs and vimentin^high^ versus vimentin^low^ macrophages (p=0.034 and p=0.114, respectively; Wilcoxon test). (d) Spatial distribution of cellular niches. Representative spatial topology of 10 K-means-clustered cellular niches (CNs) in PT and CRLM tissue sections. (e) Cell-type composition per niche. Stacked bar plots showing the proportion of 13 cell types within each CN module in PT (left) and CRLM (right) samples. (f) Enrichment and distribution of niche components. Relative intra-niche composition (left) and cross-niche distribution (right) of myCAFs, iCAFs, ECs, and vimentin^high^ macrophages in PT (top) and CRLM (bottom). (g) Spatial cell-pair adjacency counts. Abundance and proportions of direct cellular contacts among vimentin^high^ macrophages, vascular endothelial cells, and site-specific CAFs in PT and CRLM. (h) Shared-edge analysis via Voronoi tessellation. Representative Voronoi diagrams illustrating cellular physical boundary sharing among niche constituents. (i) Boundary sharing specificity between macrophages and CAFs. Quantification showing no direct spatial boundary sharing between vimentin^high^ macrophages and CAFs (p=0.295, Wilcoxon test).

To further investigate the mechanisms underlying vimentin^high^ macrophages-EC niche driven CAF phenotypes and to define the distinct contributions of each cell type, we performed Voronoi tessellation to tissue sections and quantified cell-cell boundary sharing. Our data showed that, while vimentin^high^ macrophages shared few borders with CAFs, ECs exhibited a markedly higher frequency of CAF edge-sharing. Simultaneously, vimentin^high^ macrophages were in close contact with ECs, as evidenced by the numerous shared Voronoi boundaries between them (Fig. 2h,i). These spatial arrangements suggest that ECs serve as the principal direct effector, whereas vimentin^high^ macrophages acts as the pivotal instigators within this niche, collectively orchestrating the regional specialization of CAFs across distinct tumor compartments.

### Spatial transcriptomic and mIHC validation of the vimentin^high^ macrophage-EC niche and its site-specific co-localization with CAF subtypes in CRLM

To further validate the presence and functional relevance of the vimentin^high^ macrophage-EC niche during the progression of CRLM, we integrated and analyzed spatial transcriptomic datasets (Fig. 3a), using paired PT and LM single cell transcriptomic data as reference for RCTD deconvolution (Supplementary Fig. 3a-f). In parallel, we performed mIHC staining on 38 tissue samples from 20 patients with CRLM. This analysis revealed a tumor-restricted distribution pattern: the vimentin^high^ macrophage-EC niche was consistently detected within tumor regions, yet was largely absent from adjacent normal colorectal or liver tissues (Fig. 3b).

**Figure 3.**
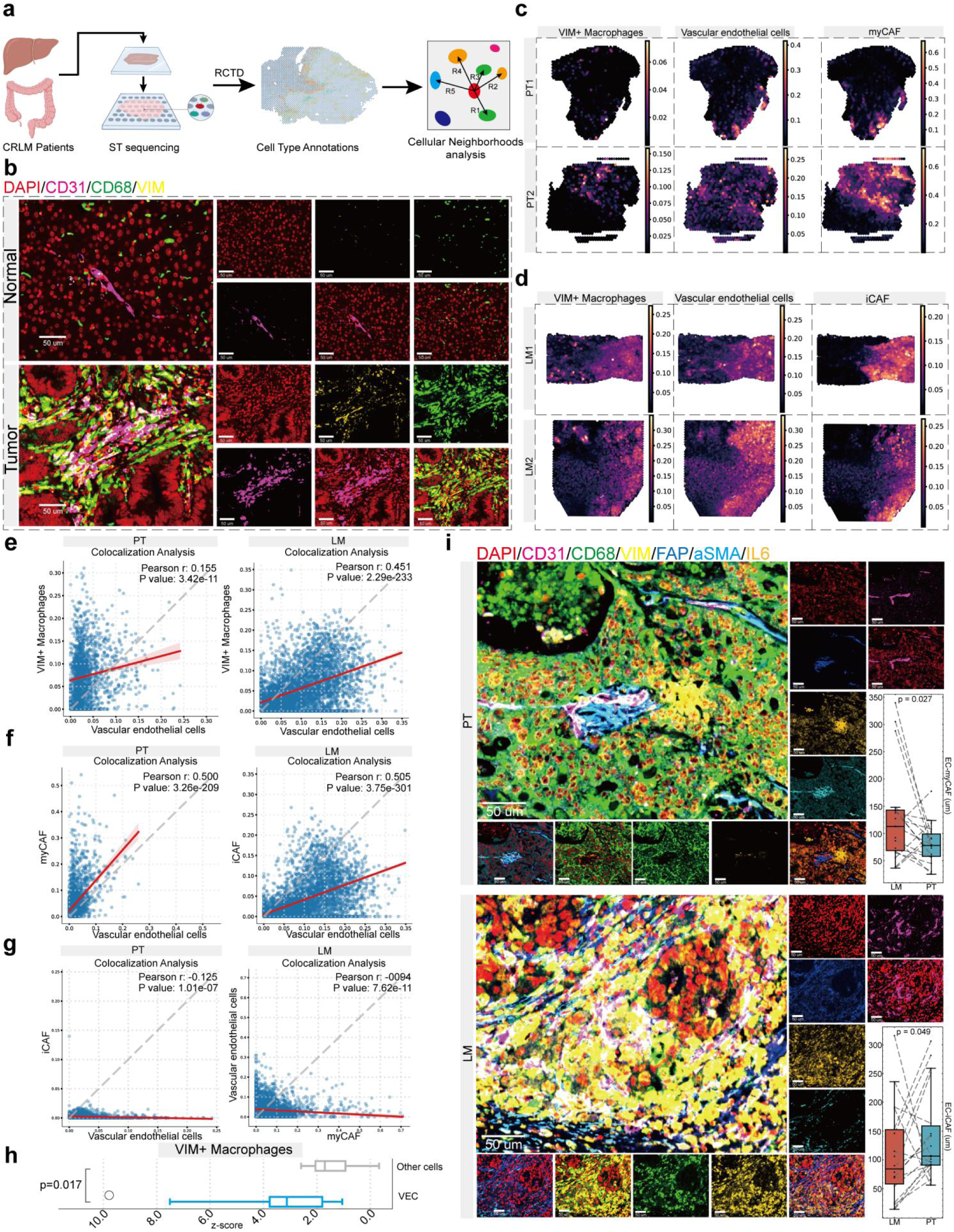
Spatial transcriptomics and multiplex imaging validate co-localization and distance gradients of the niche constituents. (a) Spatial transcriptomic deconvolution pipeline. Schematic of RCTD-based spatial transcriptomics (ST) mapping using single-cell RNA-seq reference data. (b) *In situ* detection of the vimentin^high^ macrophage–EC niche. Representative mIHC images demonstrating the restriction of ECs and vimentin^high^ macrophages signals to tumor tissue compared to normal controls (scale bars, 50 μm). (c-d) Spatial co-localization maps in ST data. Imputed spatial distribution and abundance feature maps for vimentin^high^ macrophages, vascular ECs, myCAFs in PT (c), and iCAFs in LM (d). (e-g) Quantitative spot-level co-localization analysis. Scatter plots and linear regressions showing spatial correlation between vimentin^high^ macrophages and ECs in PT/LM (e), ECs and myCAFs in PT or iCAFs in LM (f), and mutual exclusion between non-matching CAF subsets and ECs (g) (Pearson correlation, p-values indicated) (h) Permutation test for spatial co-localization. Box plots displaying significant z-score enrichment between vimentin^high^ macrophages and ECs compared to other cell pairs (p=0.017, Wilcoxon test). (i) Spatial distance quantification in multiplex immunofluorescence. Representative 7-color mIHC images (left) and spatial proximity measurements (right) showing ECs are positioned significantly closer to myCAFs in PT (p=0.027; paired Wilcoxon test; scale bars, 50 μm) and to iCAFs in LM (p=0.049; paired Wilcoxon test; scale bars, 50 μm).

Following deconvolution of each spatial transcriptomic spot to infer cell-type proportions, we performed spatial co-localization analysis on the resultant cell distributions. Consistent with our earlier observations, vimentin^high^ macrophages showed significant co-localization with ECs in both the PT and LM cohorts (Fig. 3c-e), reinforcing the robustness of this niche. Notably, in the PT group, ECs were significantly co-localized with myCAFs, whereas in the LM group, ECs preferentially associated with iCAFs (Fig. 3c,f). To more definitively ascertain the role of ECs in instructing site-specific CAF phenotypes, we examined the spatial relationships between ECs and alternative CAF subtypes. We found significant spatial exclusion between ECs and iCAFs in PT, and similarly between ECs and myCAFs in LM (Fig. 3g). Random permutation-based co-localization analysis further validated the significant association between vimentin^high^ macrophages and ECs in both cohorts (Fig. 3h).

Finally, mIHC-based spatial distance analysis further substantiated the role of this niche in orchestrating CAF heterogeneity. In the PT group, ECs within the niche were positioned significantly closer to myCAFs than to iCAFs; conversely, in the LM group, these ECs exhibited a markedly higher spatial affinity for iCAFs (Fig. 3i).

### A site-specific macrophage-EC niche reprograms CAF differentiation through divergent TGF-β and NF-κB signaling in CRLM

To elucidate the mechanisms underlying the formation of the vimentin^high^ macrophage-EC niche and its effect on CAFs, we integrated and analyzed single-cell RNA sequencing datasets from 13 patients with paired PT and LM of CRC (Fig. 4a). Batch effects were effectively mitigated using the Harmony integration framework. Following stringent quality control, a total of 373,646 high-quality cells were retained and categorized into 7 major cell lineages (Fig. 4b,c). Post-integration visualization confirmed the successful removal of batch effects (Fig. 4c). Subsequent sub-clustering of the myeloid compartment identified 7 distinct subpopulations, within which macrophages were further stratified into vimentin^high^ and vimentin^low^ subsets. Fibroblasts were similarly clustered into three subtypes: iCAFs, myCAFs, and apCAFs (Fig. 4c; Supplementary Fig. 3d-f).

**Figure 4.**
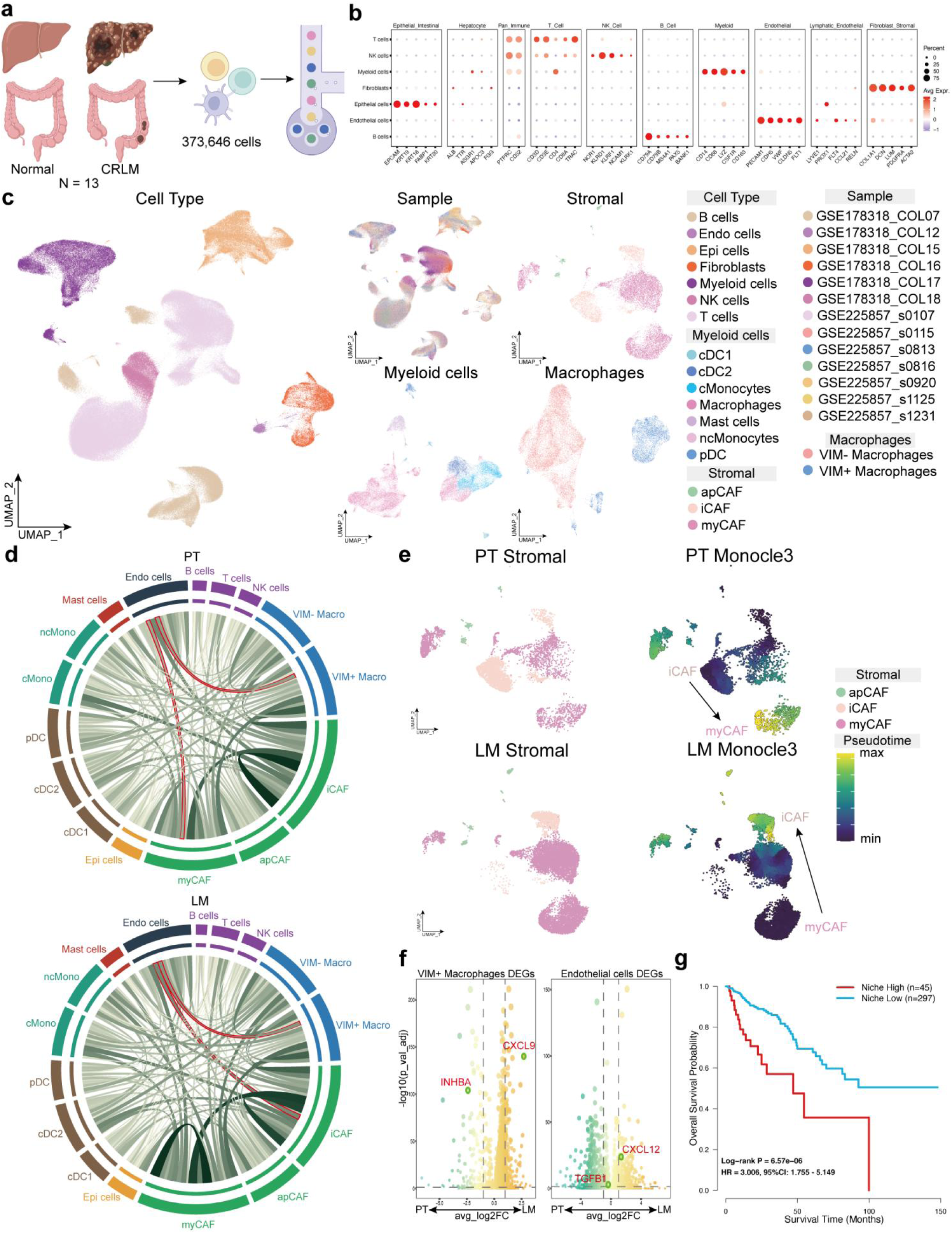
Single-cell transcriptomic profiling uncovers divergent cytokine drivers and clinical survival association. (a) Overview of paired single-cell RNA-sequencing integration. Diagram of scRNA-seq workflow encompassing paired primary CRC and liver metastases (n=13 patients, 373,646 cells). (b) Marker expression dot plot. Canonical transcript markers defining 7 main cell lineages. (c) UMAP visualization of cell populations. Dimension reduction plots displaying major cell lineages, sample distribution, sub-clustered myeloid populations, and stromal/CAF subtypes. (d) Intercellular communication networks. CellChat chord diagrams illustrating ligand-receptor interaction intensity centered on ECs, vimentin^high^ macrophages, and CAF subtypes in PT (top) and LM (bottom). (e) Pseudotime trajectory of CAF polarization. Monocle3 developmental trajectories demonstrating divergent CAF differentiation paths toward myCAFs in PT and iCAFs in LM. (f) Differential expression of niche-specific cytokines. Volcano plots highlighting significant upregulation of INHBA and TGFB1 in PT, versus CXCL9 and CXCL12 in LM across vimentin^high^ macrophages (left) and ECs (right). (g) Kaplan-Meier survival analysis. Overall survival of CRC patients stratified by vimentin^high^ macrophage–EC niche ssGSEA signature scores (p=6.57×10^−6^, HR = 3.006, 95% CI: 1.755-5.149, Log-rank test).

Cell-cell communication analysis revealed robust interactions between ECs and vimentin^high^ macrophages, as well as between ECs and both iCAFs and myCAFs, further corroborating the existence of this niche and its regulatory influence on CAF populations (Fig. 4d). Interestingly, Monocle3 trajectory analysis demonstrated that CAF differentiation bifurcated toward a myCAF phenotype in PT, whereas it preferentially shifted toward an iCAF phenotype in LM (Fig. 4e), suggesting that the niche not only recruits CAFs but also actively redirects their differentiation trajectory. To identify key cytokines driving these divergent outcomes, we performed DEG analysis on vimentin^high^ macrophages and ECs separately (Fig. 4f, Supplementary Table 5-6). INHBA secretion was significantly elevated in PT-derived vimentin^high^ macrophages, whereas CXCL9 was upregulated in their LM-derived counterparts. Concurrently, ECs in PT significantly overexpressed TGFB1, a known promoter of myCAF formation, while ECs in LM showed markedly upregulated of CXCL12, which sustains the iCAF phenotype (Fig. 4f), thus providing a mechanistic basis for the differential effects on CAF fate specification.

We next evaluated the clinical impact of this niche on patient outcomes (Fig. 4g). Survival analysis indicated that a high niche score was significantly associated with poorer prognosis; patients in the high-score group exhibited a 3.006-fold increased risk of mortality compared with the low-score group (Hazard Ratio = 3.006, 95% CI: 1.755-5.149).

The formation of distinct CAF subtypes is closely linked to their biological relevance. Integrated analyses of single-cell and bulk RNA-seq clinical cohort data revealed that myCAF generation is strongly associated with CRC progression and distant metastasis (Supplementary Fig. 4c-g), whereas iCAF formation correlates with worse immunotherapy responses (Supplementary Fig. 4h-j), underscoring the clinical significance of the vimentin^high^ macrophage-EC niche in steering CAF polarization.

The discriminatory capacity of vimentin as a functional marker for macrophage stratification remains largely unexplored. Through DEG and GO enrichment analyses, we found that vimentin^high^ and vimentin^low^ macrophages represent distinct functional states: the former are characterized by a nucleoside metabolism-related hypoxia signature, whereas the latter are enriched for antigen-presentation pathways (Fig. 5a). These divergent metabolic and transcriptional profiles establish vimentin as a reliable marker for macrophage subpopulation partitioning (Fig. 5b).

**Figure 5.**
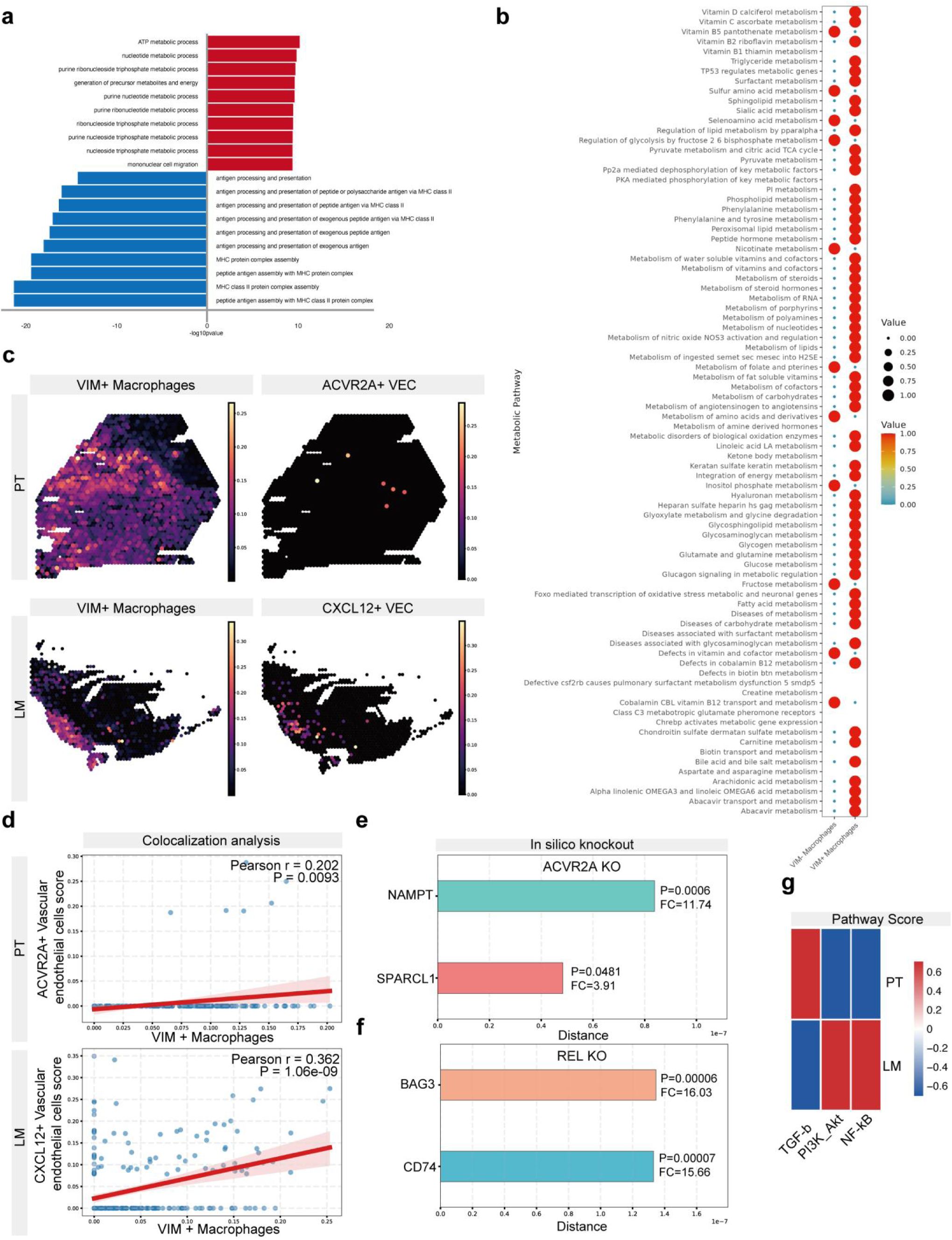
Site-specific endothelial heterogeneity and virtual knockout simulations establish the molecular cascade governing CAF polarization. (a) Functional pathway enrichment of macrophage subsets. Bar plot of GO terms showing enrichment of nucleoside/ATP metabolism in vimentin^high^ macrophages versus antigen presentation in vimentin^low^ macrophages. (b) Metabolic pathway activity dotmap. Single-cell metabolic profiling comparing vimentin^high^ and vimentin^low^ macrophage subpopulations. (c) Spatial co-localization with endothelial subsets. Spatial distribution of vimentin^high^ macrophages alongside ACVR2A+ECs in PT and CXCL12+ECs in LM. (d) Quantitative co-localization correlation. Scatter plots verifying the site-specific correlation between vimentin^high^ macrophages and ACVR2A+ ECs in PT (r=0.202, p=0.0093) or CXCL12+ECs in LM (r=0.362,p=1.06×10^−9^). (e-f) In silico knockout perturbation analysis. Predicted transcriptomic shifts and target gene alterations (SPARCL1, NAMPT, BAG3, CD74) following virtual knockout of ACVR2A in PT ECs (e) and REL (NF-κB) in LM ECs (f). (g) Pathway activity scoring. Heatmap illustrating dominant activation of TGF-β signaling in PT ECs versus PI3K-Akt and NF-κB pathways in LM EC.

Spatial analysis further revealed that vimentin^high^ macrophages preferentially co-localize with distinct EC subsets in a site-specific manner: with ACVR2A^+^ ECs in PT and with CXCL12^+^ ECs in LM, suggesting that endothelial heterogeneity mediates the niche’s effects on CAF polarization (Fig. 5c,d). Previous studies have shown that CXCL9 can indirectly trigger NF-κB-related pathways via PI3K-Akt signaling to promote CXCL12 secretion. To functionally validate these site-specific regulatory axes, we performed insilicoknockout analysis. In PT-derived ECs, silencing of ACVR2A significantly affected the expression of TGF-β pathway-related genes, including SPARCL1 and NAMPT(Fig. 5e, Supplementary Table 7). In LM-derived ECs, virtual knockout of REL, a core NF-κB family member, markedly impacted genes linked to CXCL12 regulation, including *CD74* and BAG3 (Fig. 5f, Supplementary Table 8). Consistent with these findings, pathway scoring demonstrated dominant TGF-β signaling in PT-ECs, whereas PI3K-Akt and NF-κB pathways were hyperactivated in LM-ECs (Fig. 5g).

Given that TGF-β is a master regulator of myCAF differentiation and CXCL12 is essential for iCAF maintenance, we propose a site-specific hijacking model: in PT, vimentin^high^ macrophages secrete INHBA to activate the ACVR2A/TGF-β axis in ECs, driving CAFs toward a myCAF phenotype; in LM, they secrete CXCL9 to activate the PI3K-Akt/NF-κB/CXCL12 cascade in ECs, redirecting CAFs toward an iCAF state.

### Functional validation of INHBA- and CXCL9-mediated endothelial reprogramming within the vimentin^high^ macrophage-EC niche in CRLM

To experimentally validate the proposed mechanisms, we performed invivo studies using a murine model. A highly metastatic derivative, MC38-HM, was generated from parental MC38 through six iterative cycles of intrasplenic injection (Fig. 6a). Bioluminescence imaging and gross anatomical examination both confirmed the enhanced metastatic capacity of this cell line (Fig. 6b,c). Subsequently, MC38 and MC38-HM cells were orthotopically implanted into the murine intestinal wall, and PT along with LM were harvested at experimental endpoints. While orthotopic tumors formed successfully in both groups, LM arose exclusively in the MC38-HM group (Fig. 6d). mIHC analysis of the collected specimens revealed that the presence of vimentin^high^ macrophage-EC niche in both PT and LM (Fig. 6e).

**Figure 6.**
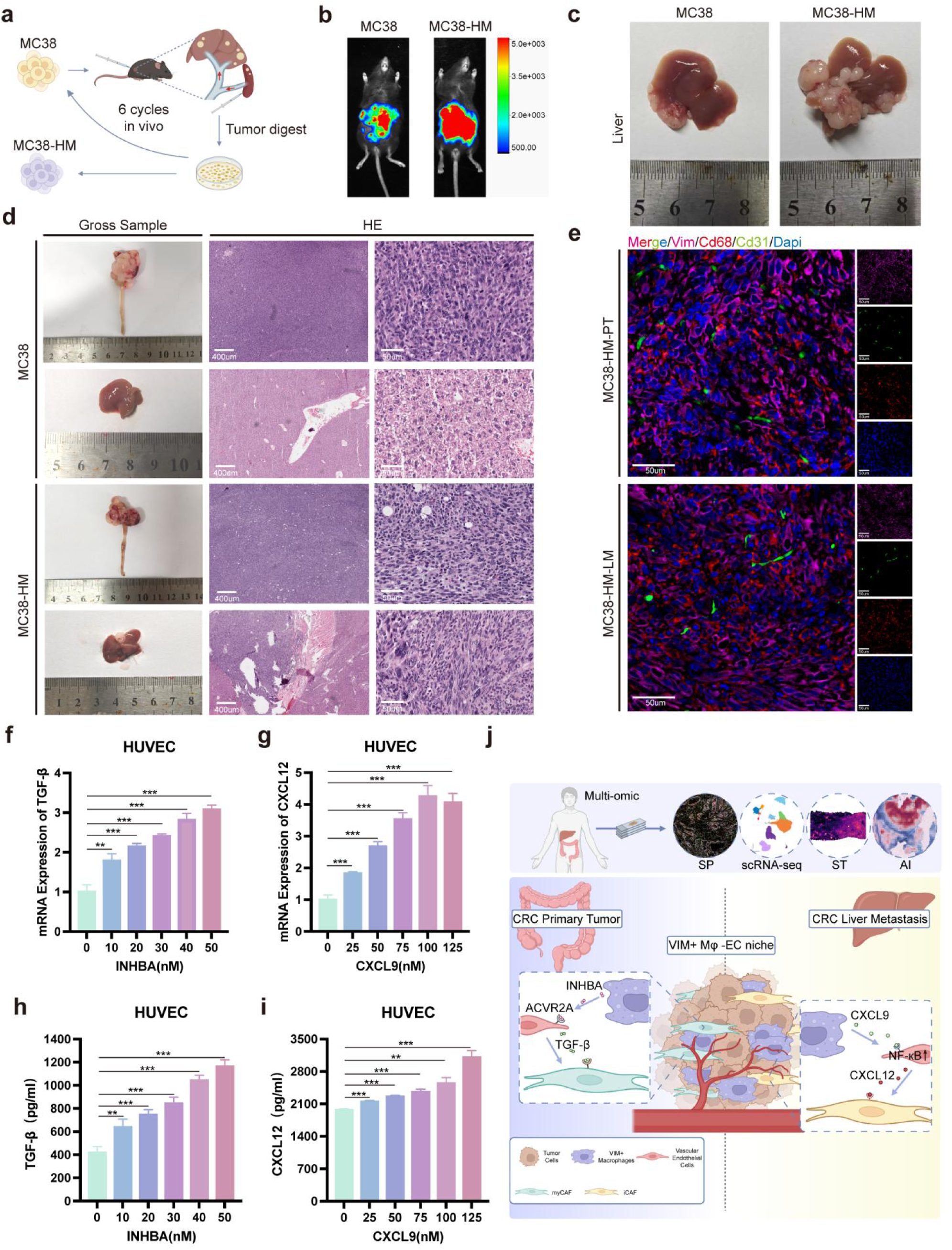
In vivo and in vitro validation confirms site-specific crosstalk driving endothelial cytokine secretion and CAF switching. (a) Establishment of the highly metastatic MC38-HM subline. Schematic of 6-cycle in vivo selection via sequential intrasplenic injection in mice. (b) Bioluminescence imaging. In vivo tracking demonstrating marked increase in hepatic metastatic burden in MC38-HM compared to parental MC38. (c) Gross liver metastasis morphology. Representative liver images from MC38 and MC38-HM injected mice. (d) Histopathological validation of orthotopic primary and metastatic lesions. Gross tissue and H&E staining of cecal primary tumors and liver metastases (scale bars, 400 μm and 50 μm). (e) *In situ* niche detection in mouse models. Multiplex immunofluorescence confirming the assembly of the Vim^+^ Cd68^+^ macrophage and Cd31^+^ EC niche in MC38-HM primary tumors (PT) and liver metastases (LM) (scale bars, 50 μm). (f-i) Recombinant cytokine stimulation of endothelial cells in vitro. Dose-dependent upregulation of TGFB1 mRNA (f) and secreted TGF-β protein (h) in HUVECs following INHBA treatment, alongside CXCL12 mRNA (g) and CXCL12 protein (i) following CXCL9 treatment (p<0.01,∗∗∗p<0.001, ANOVA). (j) Proposed mechanistic model. Graphical summary showing that in primary tumors, vimentin^high^ macrophages secrete INHBA to activate endothelial ACVR2A/TGF-β, driving myCAF polarization; in liver metastases, vimentin^high^ macrophages secrete CXCL9 to trigger endothelial PI3K-Akt/NF-κB/CXCL12 signaling, driving iCAF polarization.

To further elucidate the direct effects of INHBA and CXCL9 on ECs, we conducted invitro experiments. Treatment of HUVECs with recombinant INHBA or CXCL9 resulted in a dose-dependent increase in the mRNA levels of *TGF-β* and *CXCL12* in ECs (Fig. 6f,g). Accordingly, the protein concentrations of TGF-β and CXCL12 in the EC culture supernatants were also elevated (Fig. 6h,i), indicating that INHBA and CXCL9 independently drive EC-derived secretion of these factors.

### A deep learning framework identifies the vimentin^high^ macrophage–EC niche as a prognostic biomarker for CRC

Given the clinical significance of this niche, we explored its potential as a prognostic biomarker by developing HE2Surv, a deep learning framework integrating UNI v2 and CLAM architectures to predict survival outcomes directly from H&E-stained WSIs. UNI v2, a Vision Transformer (ViT-H)-based feature extractor, captures long-range dependencies within WSIs and exhibits strong robustness to staining and scanning variations in real-world settings. CLAM, in turn, is a weakly supervised multiple-instance learning method that incorporates attention mechanisms with clustering constraints, conferring favorable generalization performance. By combining these two approaches, HE2Surv effectively extracts histomorphological features and predicts patient prognosis. Following automated image segmentation, 560 high-quality samples were extracted for model training, while WSIs from three external centers (Zhujiang, HeBei, and GDPH) were used as a validation cohort (Fig. 7a). HE2Surv achieved high predictive accuracy in determining patient prognosis, with an AUC of 0.87762 (Fig. 7b). To ensure model transparency, we utilized attention heatmaps to visualize the histological regions most influential for its prognostic predictions (Fig. 7c). Notably, the high-attention regions identified by HE2Surv exhibited substantial spatial overlap with the vimentin^high^ macrophage-EC niche, suggesting that the model’s prognostic power derives from its capacity to recognize the spatial architecture of this specific niche. In the external validation cohorts from the three centers, HE2Surv maintained robust performance, yielding AUCs of 0.8801, 0.9167, and 0.7600, respectively, thus demonstrating favorable generalizability (Fig. 7d). Therefore, these findings position the vimentin^high^ macrophage-EC niche as a critical determinant of patient survival and highlight its potential as a novel and clinically meaningful prognostic biomarker for CRC.

**Figure 7.**
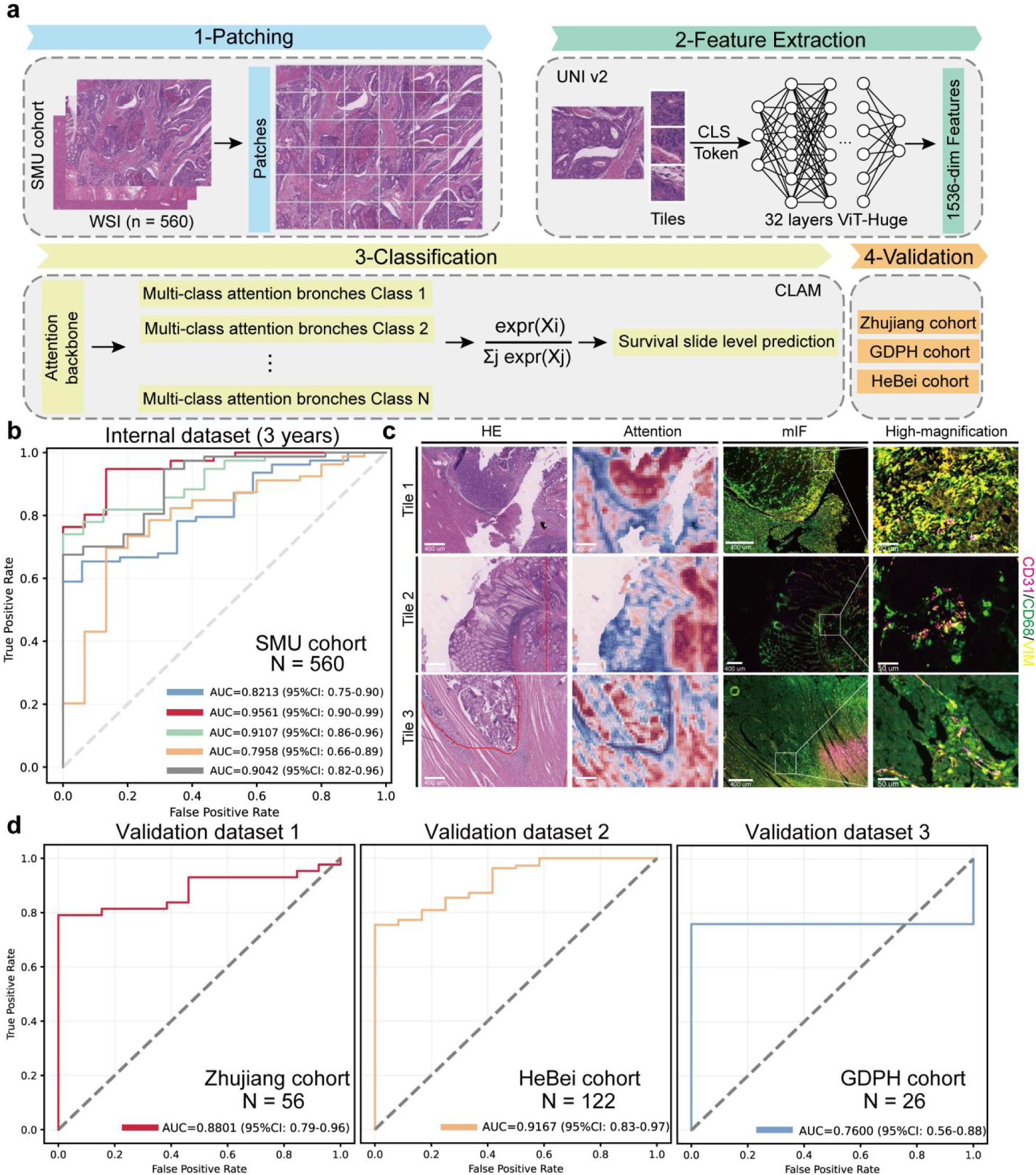
A deep-learning framework based on whole-slide histopathology captures niche architecture and predicts patient prognosis. (a) Architecture of the HE2Surv model. Schematic showing tile extraction, ViT-H feature extraction using UNI v2 (1,536-dimensional embeddings), and survival prediction using CLAM multiple instance learning. (b) Internal prognostic performance. ROC curves and AUC values of five-fold cross-validation in the primary training cohort (n=560). (c) Histopathological interpretability of HE2Surv. Attention heatmaps (middle) overlaid on H&E images (left) aligned with paired multiplex immunofluorescence (right), demonstrating that high-attention regions overlap with the spatial location of the vimentin^high^ macrophage–EC niche. (d) External validation across multi-center cohorts. ROC curves evaluating survival prediction performance in the Zhujiang (n=56, AUC = 0.8801), HeBei (n=122, AUC = 0.9167), and GDPH (n=26, AUC = 0.7600) independent validation datasets.

## DISCUSSION

Integrating ST with scRNA-seq facilitates understanding of spatial relationships among cell types^30–32^. However, direct interrogation of cell–cell interactions driving tumor immune microenvironment (TIME) remodeling and cross-sample spatial signatures remains challenging due to limited ST resolution, absence of protein-level information, and restricted sample sizes^32^. Emerging multiplex imaging platforms—including tissue-based cyclic immunofluorescence^33^, multiplexed ion beam imaging by time-of-flight (MIBI-TOF)^34^, imaging mass cytometry (IMC)^35^ and co-detection by indexing (CODEX)^36^—have enabled construction of single-cell-resolved, spatially anchored proteomic atlases across multiple cancer types^37^. These technologies permit detailed mapping of cellular architecture and reveal intricate spatial relationships fundamental to deciphering mechanisms of tumor evolution and therapeutic resistance. Nevertheless, current methodologies remain constrained by limitations in resolution, tissue applicability, and operational accessibility.

MiP-seq addresses these constraints by enabling integrated detection of DNA, RNA, proteins, and additional biomolecules at subcellular resolution, achieving synchronous acquisition of multidimensional molecular characteristics while preserving native tissue architecture^10,38^. Relative to conventional spatial transcriptomic approaches, MiP-seq not only resolves cellular heterogeneity within the TME but also precisely delineates spatial distribution patterns and intercellular interaction networks. This capability facilitates identification of critical cellular niches and their communication modes that drive tumor progression and metastatic dissemination. In this study, we integrated MiP-seq across paired primary CRC and liver metastases to define the spatial evolution of the CRC microenvironment during hepatic colonization.

Emerging evidence reported the association between tumor-associated macrophages (TAMs) density and the presence of CRLM, and identified TAMs as the key determinant of immune tolerance of liver metastasis^39,40^. Recent studies have found that a subset of macrophages expressing vimentin (vimentin^high^ macrophages) exhibits immunosuppressive effects in the hepatocellular carcinoma microenvironment, promoting tumor progression^13^. In atherosclerotic, activated macrophages upregulate vimentin expression and actively secrete vimentin into the extracellular space. This extracellular vimentin induces macrophages to release pro-inflammatory cytokines and amplifies oxidized low-density lipoprotein (oxLDL)-triggered production of TNF-α and IL-6^41^.Our study revealed that vimentin^high^ macrophages and endothelial cells form specific cellular niches in both primary colorectal cancer lesions and liver metastases. Previous researches reported macrophages accumulate abundantly within tumors, frequently establishing direct contact with ECs lining uncoated or partially coated tumor vessels^42–44^. These observations align with analyses of human cancer specimens, where elevated macrophage densities often correlate with increased angiogenic activity^45^. Collectively, these findings suggest that macrophages may exert direct proangiogenic functions in the tumor microenvironment. Mechanistic studies demonstrate that the proangiogenic activity of macrophages encompasses both secretion of classic proangiogenic factors and physical association with sprouting vessels^46^. The latter necessitates direct EC engagement with M2-like macrophages, a process regulated in part by ANG2/TIE2 and CXCL12/CXCR4 signaling axes^47,48^.Moreover, macrophage-EC interactions are bidirectional: in vitro co-culture studies demonstrate that EC monolayers can support differentiation of M2-like macrophages from myeloid progenitors^44,49^. Our study revealed the distribution characteristics and functional status of vimentin^high^ macrophages in CRC, as well as their functional interactions with ECs. However, whether there are site-specific differences in the spatiotemporal proximity between vimentin^high^ macrophages and ECs in primary lesions versus liver metastases, and their impacts on vascular remodeling, immune suppression, and metastatic colonization, remains to be further elucidated.

During malignant progression, colorectal cancer actively remodels the tumor microenvironment to foster survival and dissemination. Our analyses reveal that vimentin^high^ macrophage-endothelial cell niches in primary lesions promote differentiation of myCAFs. These myCAFs exhibit elevated α-smooth muscle actin (αSMA) and extracellular matrix (ECM)-related gene expression, conferring contractile properties and matrix deposition capacity. Through ECM remodeling and generation of pro-migratory mechanical tension, myCAFs facilitate colorectal cancer cell dissemination and metastatic progression^50,51^. Studies demonstrate that myCAF generation promotes epithelial-mesenchymal transition (EMT) and facilitates tumor metastasis. Notably, our findings reveal that vascular ECs promote myCAF formation, with spatial proximity between these populations suggesting that myCAFs may exert mechanical tension on the vascular endothelium, thereby generating physical conduits for tumor cell intravasation into the bloodstream^52^. Concurrently, TGF-β secreted by vascular endothelial cells and myCAFs may chemically compromise endothelial barrier integrity^53,54^. In liver metastases, we observed that this niche induces iCAF generation through vascular endothelial cell signaling. Vascular endothelial cells constitute the principal conduit for immune cell infiltration into tumors. iCAFs may secrete chemokines such as CXCL12 and IL-6 to recruit anti-tumor immune cells and sequester them at the tumor periphery, thereby preventing effective intratumoral penetration and cytotoxic function. Prior studies indicate that specific cancer-associated fibroblast subsets enriched at the vascular margins of colorectal cancer liver metastases correlate closely with CD8⁺ T cell exclusion^55,56^. Concurrently, establishment of colorectal cancer liver metastases requires tumor cells to transition from a “motile metastatic” to a “proliferative colonization” mode. iCAFs facilitate this process through hepatocyte growth factor (HGF) secretion, thereby promoting metastatic colonization^57,58^. Collectively, our findings demonstrate that colorectal cancer actively remodels the tumor microenvironment to drive malignant progression through vimentin^high^ macrophage-endothelial cell niche formation mediated by distinct cancer-associated fibroblast subsets. These insights highlight the therapeutic potential of targeting this cellular axis in clinical settings.

## Author Contributions

M.Z.L., B.Y.X conceived and designed the study. B.Y.X Conducted the bioinformatics analysis. M.Z.L., J.Q.W. and Z.Y.Z performed the experiments and analyzed the data. M.Z.L. and B.Y.X wrote the first draft of the manuscript. The corresponding author L.L. coordinated the overall organization, design, and writing of the article. All other authors contributed equally to the conception of the study, literature review, and editing of the manuscript and figures. All authors approved the final version of the manuscript.

## Acknowledgments

The authors thank all participants and collaborators from Zhu jiang Hospital, Fourth Affiliated Hospital of Hebei Medical University and Guangdong Provincial People’s Hospital involved in this study for their valuable contributions.

## Data Availability

The raw sequencing data reported in this study have been deposited in the Genome Sequence Archive (GSA) at the National Genomics Data Center (NGDC), China National Center for Bioinformation/Beijing Institute of Genomics, Chinese Academy of Sciences (accession No. PRJCA072405) and are publicly accessible at https://ngdc.cncb.ac.cn/gsa. The MiP-seq data for paired primary CRC and liver metastasis samples are available in the NGDC database under accession nos. OMIX019877. The original code generated for this study is accessible on the github platform at https://github.com/JavaScriptHTML5/CRC-spatial-multi-omics-analysis-MiP-seq-ST-scRNA-seq.

## Ethics Statement

The collection of clinical materials was reviewed and approved by the Institutional Committee of Southern Medical University and Nanfang Hospital (Guangzhou, China). The animal studies were approved by the Committee on the Ethics of Animal Experiments of Southern Medical University.

## REFERENCES AND NOTES

1 Bray, F. et al. Global cancer statistics 2022: GLOBOCAN estimates of incidence and mortality worldwide for 36 cancers in 185 countries. CA Cancer J Clin, doi:10.3322/caac.21834 (2024).

2 Siegel, R. L., Miller, K. D. & Jemal, A. Cancer statistics, 2020. CA Cancer J Clin 70, doi:10.3322/caac.21590 (2020).

3 Siegel, R. L. et al. Colorectal cancer statistics, 2017. CA Cancer J Clin 67, 177–193, doi:10.3322/caac.21395 (2017).

4 Cañellas-Socias, A., Sancho, E. & Batlle, E. Mechanisms of metastatic colorectal cancer. Nat Rev Gastroenterol Hepatol, doi:10.1038/s41575-024-00934-z (2024).

5 Wang, Y. et al. Liver metastasis from colorectal cancer: pathogenetic development, immune landscape of the tumour microenvironment and therapeutic approaches. J Exp Clin Cancer Res 42, 177, doi:10.1186/s13046-023-02729-7 (2023).

6 de Visser, K. E. & Joyce, J. A. The evolving tumor microenvironment: From cancer initiation to metastatic outgrowth. Cancer Cell 41, 374–403, doi:10.1016/j.ccell.2023.02.016 (2023).

7 Wang, F. et al. Single-cell and spatial transcriptome analysis reveals the cellular heterogeneity of liver metastatic colorectal cancer. Sci Adv 9, eadf5464, doi:10.1126/sciadv.adf5464 (2023).

8 Deng, Y. et al. Comprehensive single-cell atlas of colorectal neuroendocrine tumors with liver metastases: unraveling tumor microenvironment heterogeneity between primary lesions and metastases. Mol Cancer 24, 28, doi:10.1186/s12943-025-02231-y (2025).

9 Deng, Y. et al. Single-cell transcriptomic profiling reveals liver fibrosis in colorectal cancer liver metastasis. Exp Mol Med, doi:10.1038/s12276-025-01573-3 (2025).

10 Wu, X. et al. Spatial multi-omics at subcellular resolution via high-throughput in situ pairwise sequencing. Nat Biomed Eng, doi:10.1038/s41551-024-01205-7 (2024).

11 Sun, L. et al. Organization of mouse prefrontal cortex subnetwork revealed by spatial single-cell multi-omic analysis of SPIDER-Seq. Natl Sci Rev 13, nwag004, doi:10.1093/nsr/nwag004 (2026).

12 Zhang, Y. et al. CCL19-producing fibroblasts promote tertiary lymphoid structure formation enhancing anti-tumor IgG response in colorectal cancer liver metastasis. Cancer Cell 42, 1370–1385.e1379, doi:10.1016/j.ccell.2024.07.006 (2024).

13 Qiu, X. et al. Spatial single-cell protein landscape reveals vimentinhigh macrophages as immune-suppressive in the microenvironment of hepatocellular carcinoma. Nat Cancer, doi:10.1038/s43018-024-00824-y (2024).

14 Amersfoort, J., Eelen, G. & Carmeliet, P. Immunomodulation by endothelial cells - partnering up with the immune system? Nat Rev Immunol 22, 576–588, doi:10.1038/s41577-022-00694-4 (2022).

15 Asrir, A. et al. Tumor-associated high endothelial venules mediate lymphocyte entry into tumors and predict response to PD-1 plus CTLA-4 combination immunotherapy. Cancer Cell 40, 318–334.e319, doi:10.1016/j.ccell.2022.01.002 (2022).

16 Li, X. et al. Novel TCF21high pericyte subpopulation promotes colorectal cancer metastasis by remodelling perivascular matrix. Gut 72, 710–721, doi:10.1136/gutjnl-2022-327913 (2023).

17 Zeng, Z. et al. Cancer-derived exosomal miR-25-3p promotes pre-metastatic niche formation by inducing vascular permeability and angiogenesis. Nat Commun 9, 5395, doi:10.1038/s41467-018-07810-w (2018).

18 Khaliq, A. M. et al. Refining colorectal cancer classification and clinical stratification through a single-cell atlas. Genome Biol 23, 113, doi:10.1186/s13059-022-02677-z (2022).

19 Lavie, D., Ben-Shmuel, A., Erez, N. & Scherz-Shouval, R. Cancer-associated fibroblasts in the single-cell era. Nat Cancer 3, 793–807, doi:10.1038/s43018-022-00411-z (2022).

20 Elyada, E. et al. Cross-Species Single-Cell Analysis of Pancreatic Ductal Adenocarcinoma Reveals Antigen-Presenting Cancer-Associated Fibroblasts. Cancer Discov 9, 1102–1123, doi:10.1158/2159-8290.Cd-19-0094 (2019).

21 Cords, L. et al. Cancer-associated fibroblast classification in single-cell and spatial proteomics data. Nat Commun 14, 4294, doi:10.1038/s41467-023-39762-1 (2023).

22 Feng, Y. et al. Spatially organized tumor-stroma boundary determines the efficacy of immunotherapy in colorectal cancer patients. Nat Commun 15, 10259, doi:10.1038/s41467-024-54710-3 (2024).

23 Kobayashi, H. et al. The Origin and Contribution of Cancer-Associated Fibroblasts in Colorectal Carcinogenesis. Gastroenterology 162, 890–906, doi:10.1053/j.gastro.2021.11.037 (2022).

24 Koncina, E. et al. IL1R1+ cancer-associated fibroblasts drive tumor development and immunosuppression in colorectal cancer. Nat Commun 14, 4251, doi:10.1038/s41467-023-39953-w (2023).

25 Hu, S. et al. COL10A1+ fibroblasts promote colorectal cancer metastasis and M2 macrophage polarization with pan-cancer relevance. J Exp Clin Cancer Res 44, 243, doi:10.1186/s13046-025-03510-8 (2025).

26 Zhang, C. et al. Cancer-associated fibroblasts enhance colorectal cancer lymphatic metastasis via CLEC11A/LGR5-mediated WNT pathway activation. J Clin Invest 135, doi:10.1172/jci194243 (2025).

27 Li, M. et al. NEK8 kinase-mediated lactate increase impairs antitumor immunity decreasing radiotherapy sensitivity in colorectal cancer. Nat Commun, doi:10.1038/s41467-026-70657-z (2026).

28 Li, M. et al. PREX2 contributes to radiation resistance by inhibiting radiotherapy-induced tumor immunogenicity via cGAS/STING/IFNs pathway in colorectal cancer. BMC Med 22, 154, doi:10.1186/s12916-024-03375-2 (2024).

29 Liu, Z. et al. BEST: a web application for comprehensive biomarker exploration on large-scale data in solid tumors. Journal of Big Data 10, 165, doi:10.1186/s40537-023-00844-y (2023).

30 Liu, Y. M. et al. Combined Single-Cell and Spatial Transcriptomics Reveal the Metabolic Evolvement of Breast Cancer during Early Dissemination. Adv Sci (Weinh) 10, e2205395, doi:10.1002/advs.202205395 (2023).

31 Moncada, R. et al. Integrating microarray-based spatial transcriptomics and single-cell RNA-seq reveals tissue architecture in pancreatic ductal adenocarcinomas. Nat Biotechnol 38, 333–342, doi:10.1038/s41587-019-0392-8 (2020).

32 Longo, S. K., Guo, M. G., Ji, A. L. & Khavari, P. A. Integrating single-cell and spatial transcriptomics to elucidate intercellular tissue dynamics. Nat Rev Genet 22, 627–644, doi:10.1038/s41576-021-00370-8 (2021).

33 Lin, J. R. et al. Highly multiplexed immunofluorescence imaging of human tissues and tumors using t-CyCIF and conventional optical microscopes. Elife 7, doi:10.7554/eLife.31657 (2018).

34 Keren, L. et al. A Structured Tumor-Immune Microenvironment in Triple Negative Breast Cancer Revealed by Multiplexed Ion Beam Imaging. Cell 174, 1373–1387.e1319, doi:10.1016/j.cell.2018.08.039 (2018).

35 Giesen, C. et al. Highly multiplexed imaging of tumor tissues with subcellular resolution by mass cytometry. Nat Methods 11, 417–422, doi:10.1038/nmeth.2869 (2014).

36 Goltsev, Y. et al. Deep Profiling of Mouse Splenic Architecture with CODEX Multiplexed Imaging. Cell 174, 968–981.e915, doi:10.1016/j.cell.2018.07.010 (2018).

37 Risom, T. et al. Transition to invasive breast cancer is associated with progressive changes in the structure and composition of tumor stroma. Cell 185, 299–310.e218, doi:10.1016/j.cell.2021.12.023 (2022).

38 Wu, X. et al. A Guide for Spatial Omics Technologies: Innovation, Evaluation, and Application. Adv Sci (Weinh), e20806, doi:10.1002/advs.202520806 (2026).

39 Grossman, J. G. et al. Recruitment of CCR2+ tumor associated macrophage to sites of liver metastasis confers a poor prognosis in human colorectal cancer. Oncoimmunology 7, e1470729, doi:10.1080/2162402x.2018.1470729 (2018).

40 Wu, Y. et al. Spatiotemporal Immune Landscape of Colorectal Cancer Liver Metastasis at Single-Cell Level. Cancer Discov 12, 134–153, doi:10.1158/2159-8290.Cd-21-0316 (2022).

41 Kim, S. et al. Oxidized LDL induces vimentin secretion by macrophages and contributes to atherosclerotic inflammation. J Mol Med (Berl) 98, 973–983, doi:10.1007/s00109-020-01923-w (2020).

42 Squadrito, M. L. & De Palma, M. Macrophage regulation of tumor angiogenesis: implications for cancer therapy. Mol Aspects Med 32, 123–145, doi:10.1016/j.mam.2011.04.005 (2011).

43 Lin, E. Y. et al. Macrophages regulate the angiogenic switch in a mouse model of breast cancer. Cancer Res 66, 11238–11246, doi:10.1158/0008-5472.Can-06-1278 (2006).

44 Choi, Y. et al. Single-cell transcriptomics of the myeloid milieu reveals an angiogenic niche in triple-negative breast cancer. Exp Mol Med, doi:10.1038/s12276-025-01571-5 (2025).

45 Leek, R. D., Landers, R. J., Harris, A. L. & Lewis, C. E. Necrosis correlates with high vascular density and focal macrophage infiltration in invasive carcinoma of the breast. Br J Cancer 79, 991–995, doi:10.1038/sj.bjc.6690158 (1999).

46 Baer, C., Squadrito, M. L., Iruela-Arispe, M. L. & De Palma, M. Reciprocal interactions between endothelial cells and macrophages in angiogenic vascular niches. Exp Cell Res 319, 1626–1634, doi:10.1016/j.yexcr.2013.03.026 (2013).

47 Mazzieri, R. et al. Targeting the ANG2/TIE2 axis inhibits tumor growth and metastasis by impairing angiogenesis and disabling rebounds of proangiogenic myeloid cells. Cancer Cell 19, 512–526, doi:10.1016/j.ccr.2011.02.005 (2011).

48 Grunewald, M. et al. VEGF-induced adult neovascularization: recruitment, retention, and role of accessory cells. Cell 124, 175–189, doi:10.1016/j.cell.2005.10.036 (2006).

49 He, H. et al. Endothelial cells provide an instructive niche for the differentiation and functional polarization of M2-like macrophages. Blood 120, 3152–3162, doi:10.1182/blood-2012-04-422758 (2012).

50 Fang, H. et al. myCAF-derived exosomal PWAR6 accelerates CRC liver metastasis via altering glutamine availability and NK cell function in the tumor microenvironment. J Hematol Oncol 17, 126, doi:10.1186/s13045-024-01643-5 (2024).

51 Isella, C. et al. Stromal contribution to the colorectal cancer transcriptome. Nat Genet 47, 312–319, doi:10.1038/ng.3224 (2015).

52 Labernadie, A. et al. A mechanically active heterotypic E-cadherin/N-cadherin adhesion enables fibroblasts to drive cancer cell invasion. Nat Cell Biol 19, 224–237, doi:10.1038/ncb3478 (2017).

53 Pang, N. et al. Cancer-associated fibroblasts barrier breaking via TGF-β blockade paved way for docetaxel micelles delivery to treat pancreatic cancer. Int J Pharm, 124706, doi:10.1016/j.ijpharm.2024.124706 (2024).

54 Hanada, K. et al. Reduced lung metastasis in endothelial cell-specific transforming growth factor β type II receptor-deficient mice with decreased CD44 expression. iScience 27, 111502, doi:10.1016/j.isci.2024.111502 (2024).

55 Chang, M. et al. Integrated single-cell and spatial transcriptomic analysis reveals mCAF-SPP1⁺ macrophage-T cell crosstalk shaping immunosuppressive niches in colorectal cancer liver metastasis. J Transl Med, doi:10.1186/s12967-026-07978-6 (2026).

56 Sathe, A. et al. Colorectal Cancer Metastases in the Liver Establish Immunosuppressive Spatial Networking between Tumor-Associated SPP1+ Macrophages and Fibroblasts. Clin Cancer Res 29, 244–260, doi:10.1158/1078-0432.Ccr-22-2041 (2023).

57 Liao, S. et al. AFDN deficiency promotes liver tropism of metastatic colorectal cancer. Cancer Res, doi:10.1158/0008-5472.Can-23-3140 (2024).

58 Bhattacharjee, S. et al. Tumor restriction by type I collagen opposes tumor-promoting effects of cancer-associated fibroblasts. J Clin Invest 131, doi:10.1172/jci146987 (2021).

