## Supplementary Figures for "Spatial multi-omics analysis reveals vimentin^high^ macrophages–endothelial cells niche shapes CAFs heterogeneity in colorectal cancer metastasis"

**Supplementary Figure 1**

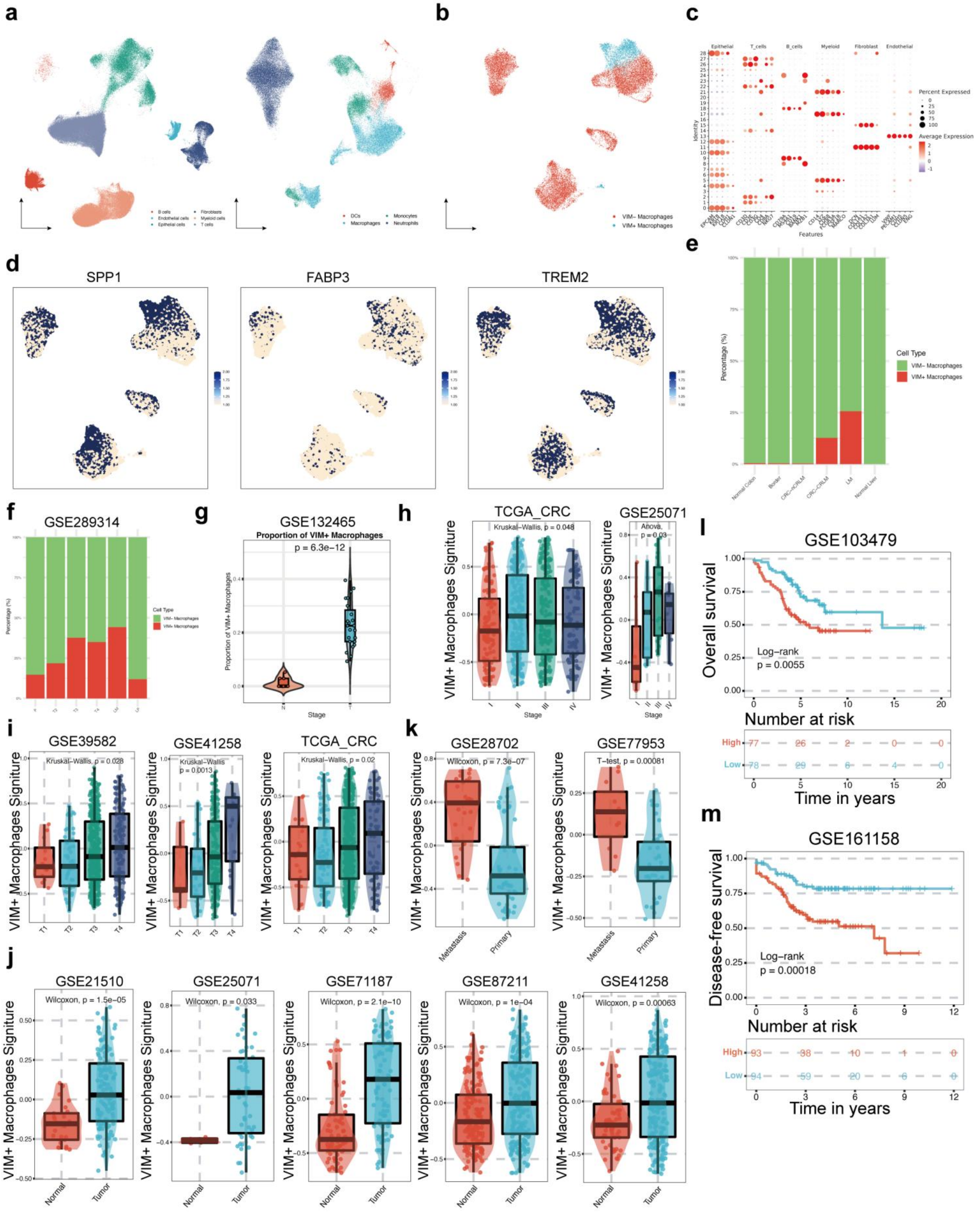

**Supplementary Figure 1. Clinical relevance and dynamic expansion of vimentin$^{high}$ macrophages during colorectal cancer progression and metastasis.**

(a) Single-cell transcriptomic atlas of colorectal cancer progression. UMAP plots showing major cell lineages across normal colon mucosa, primary tumors, invasive margins, and liver metastases.

(b) Sub-clustering of the myeloid lineage. UMAP visualization resolved into distinct myeloid subsets, emphasizing the separation of vimentin$^{high}$ and vimentin$^{low}$ macrophages.

(c) Transcriptional signature of cell lineages. Dot plot illustrating marker gene expression profiles used to define epithelial, T, B, myeloid, fibroblast, and endothelial populations.

(d) Independent expression of vimentin relative to canonical macrophage markers. Feature plots showing spatial expression of SPP1, FABP3, and TREM2 in macrophage subsets.

(e-f) Stepwise enrichment of vimentin$^{high}$ macrophages during metastatic dissemination. Stacked bar plots depicting the increasing proportion of vimentin$^{high}$ macrophages from normal tissue through primary tumors to liver metastases across integrated cohorts (e) and GSE289314 (f)

(g) Tumor-stage-dependent abundance of vimentin$^{high}$ macrophages. Box plot showing significantly higher proportions of vimentin$^{high}$ macrophages in tumor tissues compared to normal controls across disease stages (GSE132465, $p=6.3\times10^{-12}$, Kruskal-Wallis test).

(h-m) Multicenter bulk transcriptomic validation and clinical outcome association. High vimentin$^{high}$ macrophage signature scores correlate with advanced T-stage/pathological stage (h, i), tumor formation/metastasis (j, k), and significantly reduced overall survival (l; GSE103479, p=0.0055, Log-rank test) and disease-free survival (m; GSE161158, p=0.00018, Log-rank test).

**Supplementary Figure 2**

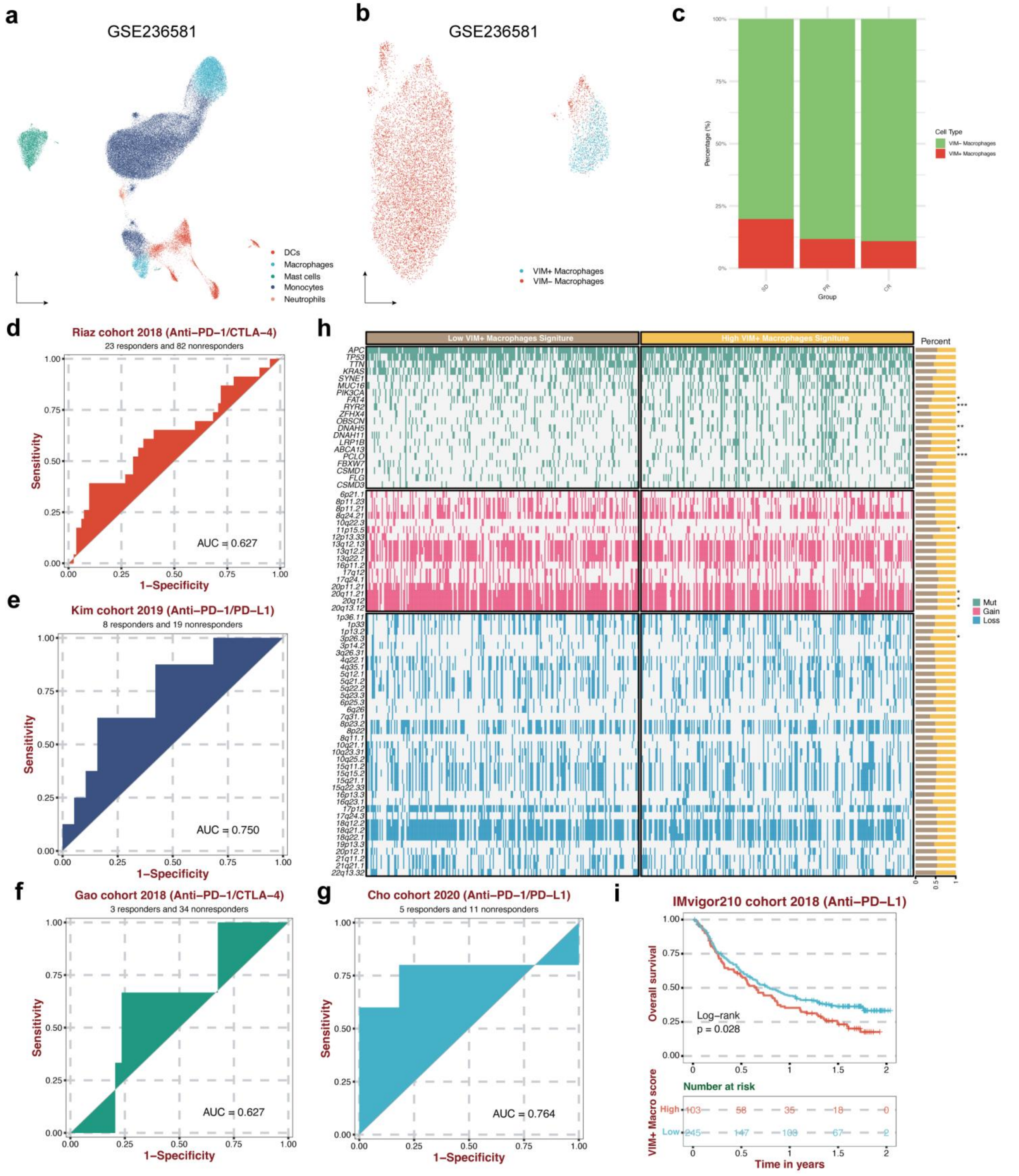


**Supplementary Figure 2. Predictive capacity of the vimentin$^{high}$ macrophage signature for immunotherapy response in CRC.**

(a-b) Single-cell profiling of CRC patients undergoing immunotherapy. UMAP visualization of myeloid subpopulations (a) and sub-clustered macrophage subsets (b) from patients receiving immune checkpoint blockade (GSE236581).

(a) Macrophage subset dynamics across treatment response groups. Proportion of vimentin$^{high}$ macrophages in patients exhibiting stable disease (SD), partial response (PR), and complete response (CR), showing a progressive decline with therapeutic response.

(d-g) Clinical prediction of immunotherapy response across independent cohorts. Receiver operating characteristic (ROC) curves evaluating the power of the vimentin$^{high}$ macrophage signature to predict response to Anti-PD-1/CTLA-4 or Anti-PD-1/PD-L1 therapies in the Riaz cohort (d, AUC = 0.627), Kim cohort (e, AUC = 0.750), Gao cohort (f, AUC = 0.627), and Cho cohort (g, AUC = 0.764).

(h) Genomic landscape and copy-number alterations. Heatmap comparing somatic mutation patterns and copy-number variation (CNV) gain/loss distributions between low and high vimentin$^{high}$ macrophage signature groups.

(i) Prognostic value of the vimentin$^{high}$ macrophage signature for overall survival. Kaplan-Meier analysis evaluating the association between the vimentin$^{high}$ macrophage signature and overall survival in the IMvigor210 cohort (2018) receiving anti-PD-L1 therapy. Patients were dichotomized into high (red) and low (blue) signature groups. High signature scores were significantly associated with worse overall survival (Log-rank $p = 0.028$).

**Supplementary Figure 3**

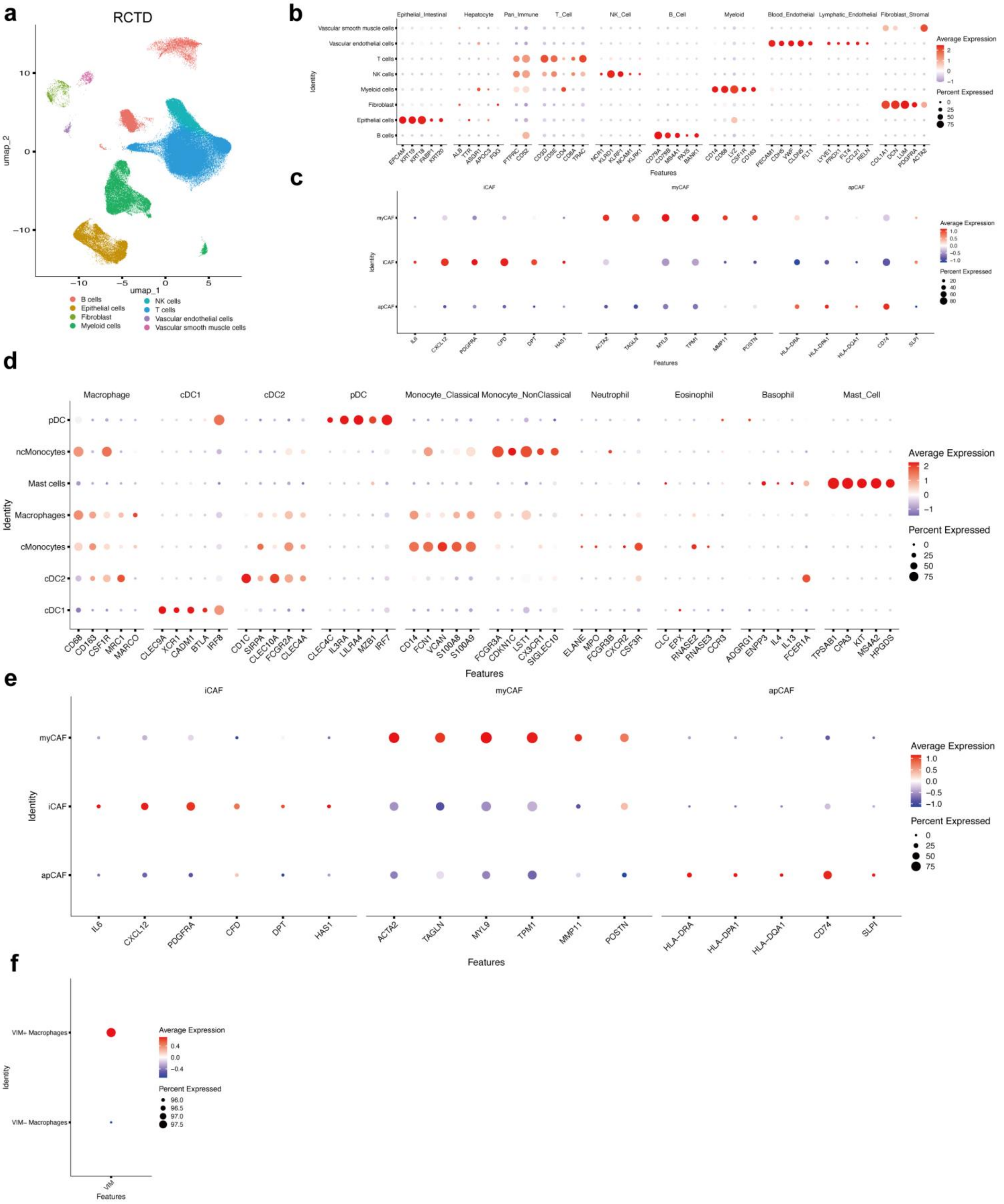


**Supplementary Figure 3. Deconvolution references and specific expression signatures for spatial and single-cell transcriptomic mapping.**

(a) Single-cell reference atlas for RCTD deconvolution. UMAP plot of the reference scRNA-seq dataset (GSE178318) used for cell-type decomposition of Visium spatial transcriptomics spots.

(b) Expression profile of major reference cell types. Dot plot showing canonical gene markers for 8 cell types included in the spatial deconvolution reference.

(c) Marker gene expression across CAF subpopulations. Dot plot displaying expression of representative markers for iCAFs (IL6, CXCL12), myCAFs (ACTA2, TAGLN), and apCAFs (HLA-DRA, CD74).

(d-f) Detailed transcriptomic signatures of sub-clustered populations. Dot plots illustrating marker gene expression profiles across 7 myeloid cell subpopulations (d), 3 functional CAF states (e), and macrophage sub-lineages defined by VIM expression (f)

**Supplementary Figure 4**

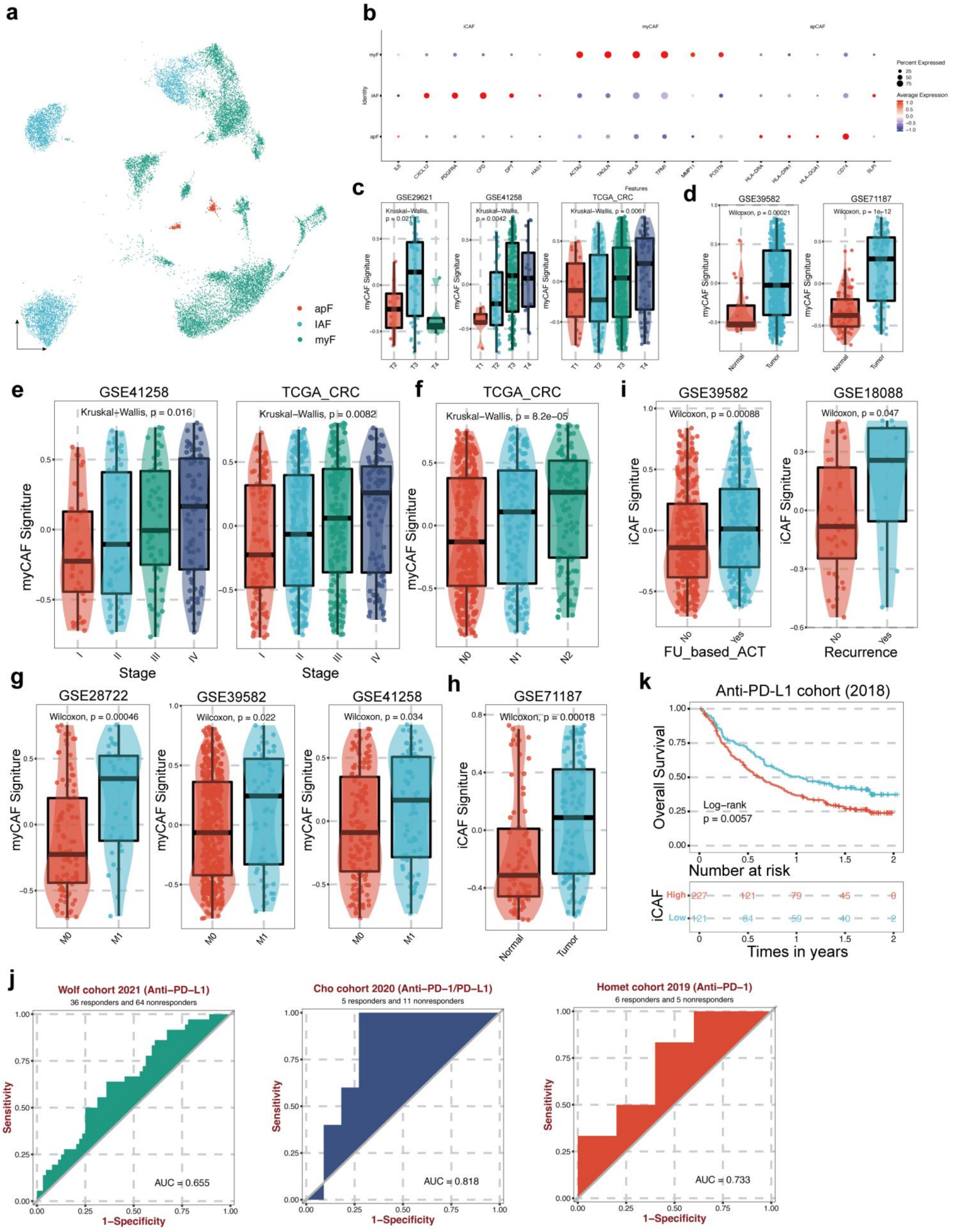


**Supplementary Figure 4. Biological significance and clinical impact of divergent CAF subsets in colorectal cancer.**

(a) UMAP visualization of fibroblast sub-clusters. Identification of myofibroblastic CAFs (myCAF), inflammatory CAFs (iCAF), and antigen-presenting CAFs (apCAF).

(b) Transcriptomic profiling of CAF subsets. Dot plot confirming subset-specific markers across myCAF, iCAF, and apCAF subpopulations.

(c-f) Correlation between myCAF abundance and clinical tumor burden. High myCAF signature scores significantly associate with advanced T-stage (c), tumor fomration (d), overall stage progression (e), and lymph node involvement (f; N-stage, $p=8.2\times10^{-5}$, Kruskal-Wallis test) across public cohorts (GSE29621, GSE41258, TCGA_CRC).

(g) Elevated myCAF signature in metastatic lesions. Increased myCAF scores in primary tumors with distant metastasis (M1) versus non-metastatic tumors (M0) across multiple datasets.

(h-j) Association of iCAFs with tumor recurrence and immunotherapy resistance. iCAF signature scores are significantly upregulated in tumor tissues versus normal controls (h) and in patients experiencing chemotherapy relapse (i). High iCAF scores predict poor response to immunotherapy in independent clinical trials (j; Wolf, Cho, and Homet cohorts).

(k) Impact of iCAF signature on overall survival. Kaplan-Meier survival curves demonstrating that high iCAF score correlates with significantly reduced overall survival in anti-PD-L1 treated patients ($p=0.0057$, Log-rank test).
